# Nanoscale architecture of a VAP-A-OSBP tethering complex at membrane contact site

**DOI:** 10.1101/2020.10.13.337493

**Authors:** Eugenio de la Mora, Manuela Dezi, Aurélie Di Cicco, Joëlle Bigay, Romain Gautier, John Manzi, Joël Polidori, Daniel Castaño Díez, Bruno Mesmin, Bruno Antonny, Daniel Lévy

## Abstract

Membrane contact sites (MCS) are subcellular regions where two organelles appose their membranes to exchange small molecules, including lipids. Structural information on how proteins form MCS is scarce. We designed an in vitro MCS with two membranes and a pair of tethering proteins suitable for cryo-tomography analysis. It includes VAP-A, an ER transmembrane protein interacting with a myriad of cytosolic proteins, and oxysterol-binding protein (OSBP), a lipid transfer protein that transports cholesterol from the ER to the trans Golgi network. We show that VAP-A is a highly flexible protein, allowing formation of MCS of variable intermembrane distance. The tethering part of OSBP contains a central, dimeric, and helical T-shape region. We propose that the molecular flexibility of VAP-A enables the recruitment of partners of different sizes within MCS of adjustable thickness, whereas the T geometry of the OSBP dimer facilitates the movement of the two lipid-transfer domains between membranes.

## INTRODUCTION

Membrane contact sites (MCS) are subcellular regions where two organelles associate to carry out non-vesicular communication (Alli-Balogun et al., 2019; Helle et al., 2013; Hoffmann and Kukulski, 2017; Prinz, 2014; Scorrano et al., 2019; Wu et al., 2018). MCS involve almost every organelle, are present in all tissues, play important roles in lipid exchange, calcium signaling, organelle fission, inheritance, and autophagy, and are implicated in metabolic diseases.

In MCS, the two facing membranes are closely apposed, typically 15–30 nm apart, over distances up to micrometers. Consequently, MCS are highly confined spaces but their molecular organization is poorly understood, notably with regards to protein stoichiometry, density, orientation and dynamics. Proteins involved in membrane tethering display a wide variety of organizations: single or multiple polypeptide chains, complexes of two proteins, each of them associated to a different organelle, or multimeric assemblies such as the ERMES complex between the ER and mitochondria (Scorrano et al., 2019).

To date, no complete structure of tethers involved in MCS formation has been solved. In general, structural information is limited to protein domains such as those involved in lipid transfer or organelle targeting (Schauder et al., 2014). Because these domains are often connected by disordered linkers, this leaves numerous possibilities for how the full-length proteins orient and move between the two facing membranes. Recently, a 3D model of an ~160-kD N-terminal fragment of the lipid transport protein VPS13 has been reported, revealing an ~160-Å long channel that enables lipid flux between organelles (Li et al., 2020). However, all structures have been determined in the absence of membranes, resulting in an incomplete view of MCS.

Thanks to advances in cryo-electron microscopy (cryo-EM), in particular in situ cryo-tomography (cryo-ET), first images of MCS at medium resolution have been obtained. Synaptotagmins and their yeast orthologs tricalbins are rod-shape 15-20 nm long structures that bridge the ER and the PM (Collado et al., 2019; Fernández-Busnadiego et al., 2015; Hoffmann and Miller).

Here we address the general question of the formation of MCS at molecular scale by studying a model MCS formed by VAP-A and OSBP. VAP-A and its homolog VAP-B are transmembrane ER proteins that interact with ≈ 100 cytosolic partners. They do so through a common mechanism: the cytosolic Major Sperm Protein (MSP) domain of VAP-A/B recognizes FFAT (two phenylalanine in an acidic track) or FFAT-like motifs in partner proteins (Huttlin et al., 2015; Lev et al., 2008; Murphy and Levine, 2015). The large spectrum of VAP-A interactants and their involvement in many MCS make the structure of VAP-A at MCS an important open question.

OSBP and OSBP-related proteins (ORPs) constitute a large family of lipid transfer proteins (LTPs). Several ORPs transport specific lipids in a directional manner owing to the counter exchange and hydrolysis of the phosphoinositide PI4P. OSBP drives cholesterol/PI4P exchange at this contact site and is the target of several anticancer and antiviral compounds (Burgett et al., 2011), pointing to its key role in cellular homeostasis.

In this work, we designed an in vitro system adapted for cryo-EM and cryo-ET analysis of MCS formed by VAP-A and either OSBP or a shorter construct, N-PH-FFAT, containing OSBP tethering determinants. By sub-tomogram averaging, we obtained 3D models of VAP-A and N-PH-FFAT between facing membranes, which reveal the organization of membrane tethering.

## RESULTS

We studied the structure of minimal membrane contact sites formed by two purified proteins, VAP-A and OSBP. The domain organization of these proteins is schematized in Figure 1A.

**Figure 1:**
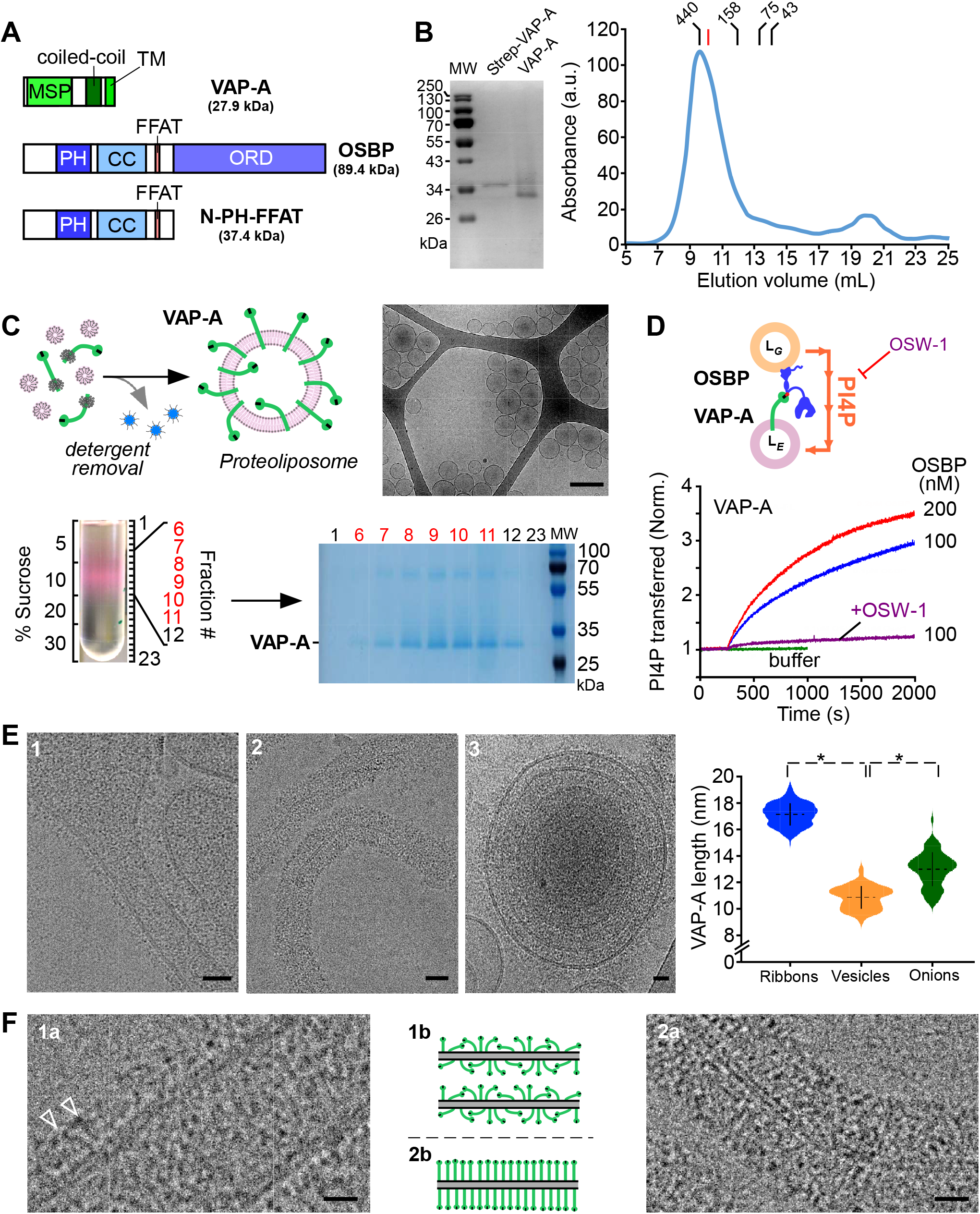
VAP-A extends from the membrane with increasing protein density. (A) Domain organization of VAP-A, of OSBP and of the N-PH-FFAT construct. (B) Purification of full length VAP-A with its transmembrane domain after solubilization in DDM. SDS-PAGE analysis of purified strepII-TEV-tag VAP-A before and after TEV proteolysis. Size-exclusion chromatography of VAP-A. The red marker is BmrA, a 130 kDa membrane protein solubilized in DDM. (C) The scheme describes the principle of proteoliposome formation by detergent removal. Lipid/protein molar ratio (LPR) before detergent removal was adjusted to change the amount of VAP-A in proteoliposomes. Floatation of VAP-A proteoliposomes doped with a fluorescent lipid. SDS-PAGE analysis shows that VAP-A is incorporated in a single population of vesicles. Cryo-EM images of VAP-A proteoliposomes. Bar = 250 nm. (D) The scheme describes the principle of PI4P translocation between Golgi-lipid vesicles and VAP-A proteoliposomes as catalyzed by OSBP. Real-time measurement of PI(4)P transfer in the presence of VAP-A proteoliposomes at LPR 4000 mol/mol. NBD-PH (3 μM) was mixed with Golgi-like liposomes (250 μM lipids containg 4 mol % PI(4)P and 2 mol % Rho-PE) and VAP-A proteoliposomes (250 μM lipids, 0.17 nM VapA). At t = 300 sec, OSBP was added at 100 or 200 nM as indicated. OSW1 (1μM) an inhibitor of OSBP was used to inhibit PI4P transfer. (E) Reconstitution of VAP-A at high protein density (LPR 70 mol/mol) leading to the formation of deformed vesicles (1), ribbons (2), onions (3). Violin plots showing the length distribution of the extramembrane region of VAP-A. The plot shows the complete distribution of values analyzed. The horizontal and vertical lines show the average value and the standard deviation, respectively. * indicates p < 0.01 by unpaired t test. 50 vesicles (n=78 measurements), 45 ribbons (n= 421 measurements) and 6 onions (n=78 measurements) were analyzed. Bar = 25 nm. (Figure S1). (F) Close-up views of regions 1a, 2a where electron densities of VAP-A are visible and schematic representation of protein concentration and orientation in the membrane in deformed vesicles 1b, 2b (tubular region of E1) and ribbons (E2). Some individual proteins are resolved (white arrow heads). The extramembrane region of VAP-A is visible and extends at increasing distances from the membrane. Bars = 10 nm (E, F).

VAP-A is inserted in the ER membrane using a single C-terminal transmembrane helix (TM) and exposes its N-terminal globular domain of the Major Sperm Protein family (MSP domain) to the cytoplasm. The MSP, which recognizes proteins that contain FFAT or FFAT-like motifs, is separated from the TM by a predicted coiled coil region (CC) (Murphy and Levine, 2015). Together with the TM, the CC promotes VAP-A dimerisation (Kim et al., 2010).

OSBP contains five functional regions: an N-terminal intrinsically disordered region, a PH domain, a putative central coil-coiled domain, a FFAT motif, and C-terminal OSBP related domain (ORD), which exchanges cholesterol for PI4P (Mesmin et al., 2013; Ridgway et al., 1992). OSBP bridges ER and TGN *via* its FFAT motif, which recognizes VAP-A, and *via* its PH domain, which recognizes Arf1-GTP and/or PI4P present in the Golgi membrane. The disordered N-terminus of OSBP limits the density of proteins at MCS and controls OSBP orientation. A simplified N-PH-FFAT construct of OSBP, lacking the ORD, recapitulates the tethering function of OSBP (Jamecna et al., 2019; Mesmin et al., 2013).

### Purification/biochemical and functional characterization of full-length VAP-A

In this study, we used full-length OSBP or N-PH-FFAT to reconstitute MCS and analyze their architecture by cryo-EM and cryo-ET. To better mimic the cellular conditions, we used full length VAP-A with its TM inserted into the liposome bilayer.

We expressed full length VAP-A in *E. coli* and purified it in a micellar form using the mild detergent n-dodecyl-β-D-maltoside (DDM) (Figure 1B). Size exclusion chromatography showed a single peak at MW ~ 420 kDa. As observed for membrane proteins, the apparent MW was higher than that expected for a VAP-A monomer (27.9 kDa) or dimer (54.8 kDa) due to the presence of the DDM micelle. However, the elution peak of VAP-A also preceded that of BmrA, a membrane protein of 130 kDa solubilized in DDM, suggesting that VAP-A had an oligomeric form larger than a dimer or a non-globular form.

We reconstituted VAP-A in liposomes by adding phosphatidylcholine (PC) and phosphatidylserine (PS) to the VAP-A-DDM micelles, followed by detergent removal (Figure 1C). We chose a high lipid/protein ratio (LPR ≈ 1400 mol/mol) to ensure that proteins were present at low density in the membrane. CryoEM showed that the proteoliposomes were spherical, unilamellar, and displayed a diameter ranging from 40 to 200 nm. After flotation on a sucrose gradient, the proteoliposomes were recovered in a single low sucrose density band, suggesting a homogeneous population (Figure 1C, lanes 6 to 11). No protein aggregates were found at the bottom of the gradient (lane 23 in Figure 1C).

To test if VAP-A in proteoliposomes was able to functionally interact with OSBP (Figure 1D), we used an assay that follows the transfer of PI4P from Golgi-like to ER-like liposomes in real time (Mesmin et al., 2013). We incubated the VAP-A proteoliposomes with Golgi-like liposomes containing 4% PI(4)P and Rhodamine lipid (Rho-PE), and used the probe NBD-PH as a fluorescent reporter of the membrane distribution of PI(4)P. At the beginning of the experiment, the fluorescence of NBD-PH was quenched by Rhodamine lipid (Rho-PE) since both PIP4 and Rho-PE were present in the same liposomes. The addition of OSBP triggered a large increase in the fluorescence of NBD-PH due to the transfer of PI(4)P to the VAP-A proteoliposomes, which did not contain Rho-PE (Figure 1D). PI4P transfer rate increased with the amount of OSBP and was inhibited by OSW1, a specific OSBP inhibitor.

These results show that full length VAP-A purified in DDM is well folded, can be incorporated in lipid membranes and is functional.

### VAP-A extends at increasing distances from the membrane with its concentration

We reconstituted VAP-A at LPRs ranging from 2800 to 50 mol/mol. We expected that increasing VAP-A density could favor protein-protein interaction in the membrane plane. We visualized the reconstituted proteoliposomes by cryo-EM. When the starting mixture was at LPR 70 mol/mol, i.e a high protein density, the reconstituted proteoliposomes were heterogeneous in shape and size. We observed (i) spherical vesicles, (ii) non-spherical vesicles displaying angular shapes or tubulation, (iii) fragments of open membranes resembling ribbons with proteins on both side of the bilayer, and (iv) multilayered vesicles (Figures 1E, S1A). When the concentration of VAP-A was reduced to LPR 350 mol/mol, the fraction of spherical vesicles increased at the expense of ribbons and deformed vesicles (Figure S1A–S1D). At very low protein density (LPR = 700-1400 mol/mol), all proteoliposomes were spherical or slightly deformed. Thus, vesicle morphology depended on the concentration of VAP-A in the membranes (Figures S1E-L). High protein density shapes the membrane probably by crowding effect and/or lateral protein-protein interactions, as shown for other transmembrane proteins reconstituted at high density for electron crystallography and AFM (for reviews see (Hasler et al., 1998; Rigaud et al., 2000).

Even though VAP-A has a small size for cryo-EM, electron densities of VAP-A extending out of the membrane of vesicles were clearly visible. We measured the length of the extramembrane domain of VAP-A in different types of vesicles after reconstitution at LPR 70 mol/mol using the well-resolved electron density of the external lipid leaflet. VAP-A extended 10±2 nm (n=284 measurements, 50 vesicles) from the outer lipid leaflet of deformed vesicles. In some cases, individual proteins with an elongated shape were identified (Figure 1F, 1a–1b, white arrows). At the protein tip, dark dots likely corresponding to the MSP domain were followed by a region toward the membrane without defined structure. Some dark dots were also found closer to the membrane suggesting that VAP-A and/or its MSP domains adopted various orientations with regard to the membrane plane. In the case of ribbons (Figure 1F, 2a–2b), VAP-A appeared more compact and perpendicular to the lipid bilayer, extending up to 17 ± 2 nm (n= 422 measurements, 50 ribbons). This elongated shape is consistent with the high apparent molecular weight as observed by size-exclusion chromatography (Figure 1A).

**Figure 2.**
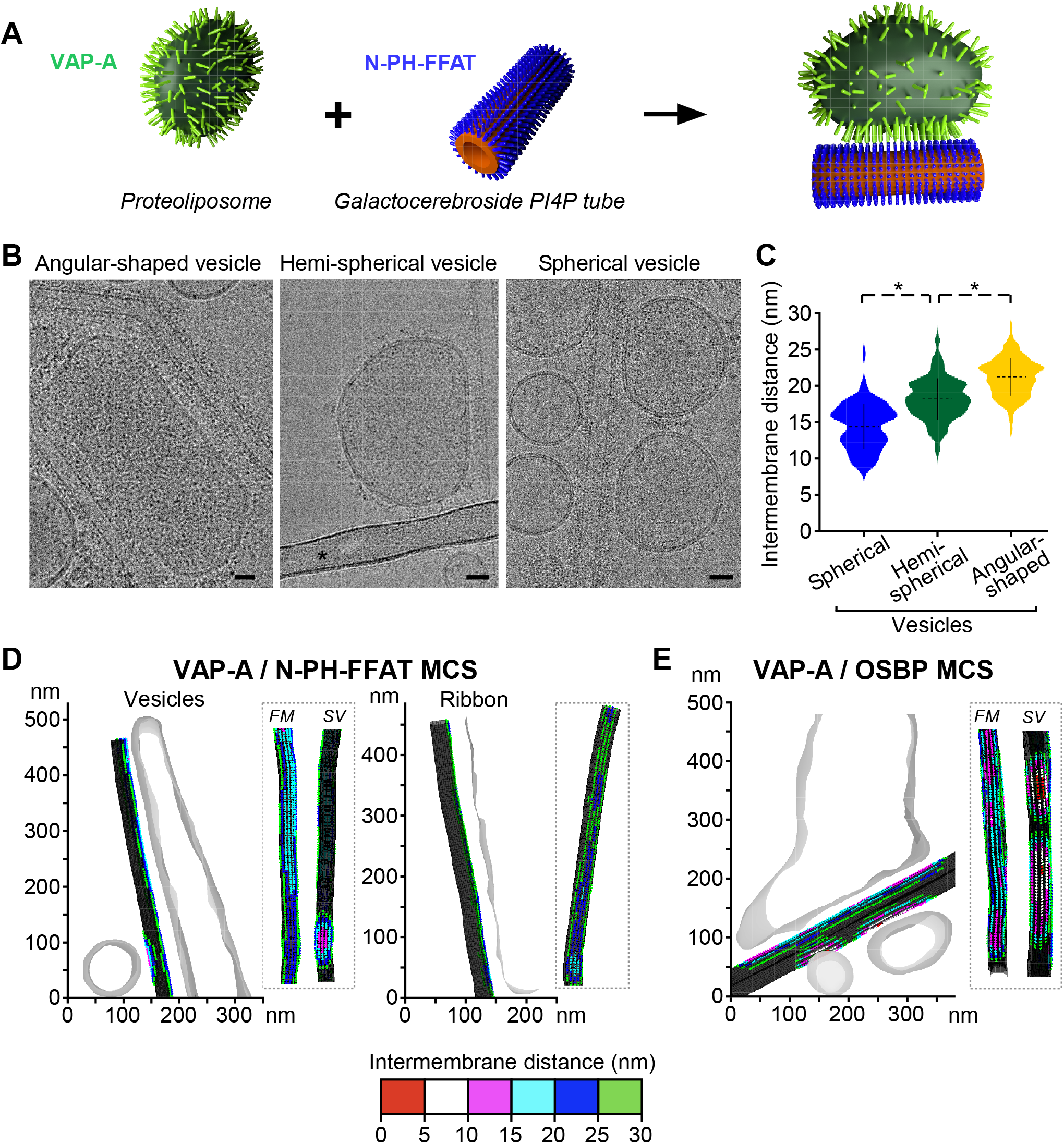
VAP-A density tunes the separation between facing membranes in reconstituted MCS. (A) Scheme of the reconstitution of MCS between VAP-A proteoliposomes and N-PH-FFAT (or OSBP) bound to galactocerebroside tubes doped with PI4P. (Figure S2A) (B) Representative images of the different types of contacts made between VAP-A proteoliposomes (LPR 70 mol/mol) and N-PH-FFAT bound to tubes. The bar labeled with * in the central panel is a carbon grid bar. Bars = 25 nm. (Figure S2B) (C) Violin plots of the distance between facing membranes in reconstituted MCS (LPR 70 mol/mol). The plots show the complete distribution of values analyzed. The horizontal and vertical lines show the average value and the standard deviation, respectively. * indicates p < 0.01 by unpaired t test. A total of 90, 122 and 208 distances of contact made between tubes and spherical, hemi-spherical and angular-shaped vesicles were analyzed. (D) 3D reconstruction of VAP-A/N-PH-FFAT and (E) VAP-A/OSBP MCS from cryo-tomograms. Distances between angular shape and spherical vesicles and tubes and ribbons and tubes are depicted in color codes on tubes. *FM* indicates the tube side in contact with a flattened region of large vesicles and *SV* the tube side in contact with small spherical vesicles. (Movies S1-S3).

The multilayered vesicles were made of concentric membrane layers separated by protein densities, suggesting that they were held together by VAP-A homotethers (Figure 1E3). This organization was reminiscent of that of *Ascaris suum* MSP dimers, which assembled antiparallel due to 3D crystal packing (Bullock et al., 1996). VAP-B homotether existence has been suggested to explain stacked ER cisternae phenotype in Nir2/VAP-B-expressing cells (Amarilio et al., 2005). The separation between facing membranes in multilayered vesicles was 22-28 nm, suggesting that VAP-A molecules extend 11 to 14 nm from the membrane.

In summary, VAP-A extends at variable distances from the membrane, adopting either a compact and tilted conformation at low surface density or an extended conformation at high surface density, suggesting a highly flexible molecule.

### Formation of membrane contact sites with VAP-A and OSBP

We next studied how VAP-A binds partners and forms a membrane contact site. We used either full-length OSBP or its N-PH-FFAT construct, which contains the membrane determinants for bridging VAP-A to PI4P-containing membranes.

Because VAP-A and N-PH-FFAT are small (MW = 55.8 kDa and 74.8 kDa as dimers, respectively), we suspected that the putative VAP-A-N/PH-FFAT complex of 130.6 kDa might be difficult to recognize within a contact zone where proteins are densely packed. We thus designed an *in vitro* system that could allow us to assign the membrane system to which each protein was bound. Specifically, we mixed two morphologically different membranes: VAP-A was incorporated into ER-like liposomes, while N-PH-FFAT was bound to lipid tubes made of galactocerebroside (GalCer) containing PI(4)P (Figure 2A). The lipid tubes were also doped with PC and PS to increase membrane fluidity (Wilson-Kubalek et al., 1998). They were of constant diameter of 27 nm and showed variable lengths of several hundreds of nanometers (Figure S2A). N-PH-FFAT or OSBP were bound at high concentration to the tubes as shown by the protein densities covering the external lipid leaflet and extending 5-6 nm from the membranes (Figure S2A, white arrow heads).

We incubated VAP-A proteoliposomes with PI4P tubes decorated with N-PH-FFAT and froze the mixture for cryo-EM observation. After 2 min incubation, we observed massive aggregation of vesicles and tubes suggesting fast tethering of the two membranes (Figures S2B, S2D). To reduce aggregation, we added sequentially both components directly onto the cryo-EM grid, before freezing within less than 30 seconds (Figures S2C, S2E). Contact areas were recognizable at low magnification with remodeled vesicles in close proximity of tubes (Figure S2E white arrow). This protocol resulted in an ice layer that was thin enough for cryo-EM or cryo-ET and was therefore selected for further analysis of MCS architecture and formation.

### The MCS inter membrane distance depends on the local concentration of VAP-A

VAP-A is present along the ER and concentrates in MCS, e.g. at the ER-Golgi or ER-plasma membrane interface where FFAT-containing proteins such as OSBP or Kv2.2 channel, respectively, are present (Mesmin et al., 2013;Johnson et al., 2018). However, the number of VAP-A molecules engaged in MCS is not known. We analyzed how MCS assembled when the concentration of VAP-A in ER-like membrane varied.

We reconstituted VAP-A at a low LPR ratio (70 mol/mol) to obtain a mix of spherical vesicles, of deformed vesicles and of membrane ribbons. After incubation with N-PH-FFAT tubes, different types of contact areas were observed depending on the morphology of the VAP-A-containing bilayer. We identified (i) angular-shaped proteoliposomes with high protein density on the surface and with membrane lying along the tubes, (ii) hemi-spherical vesicles with deformations restricted to a part of the vesicles, and (iii) spherical liposomes in tangential contact with the tubes (Figures 2B, 2D). The separation between the facing membranes was not constant and depended on the class of contacting vesicles. Measurements from 2D images showed an intermembrane distance between the tubes and the VAP-A-containing membranes of 15 nm ± 5 nm for the spherical vesicles to 20 ± 5 nm for the highly deformed vesicles (Figure 2C). Larger distances up to 30 nm were measured in cryo-tomograms of VAP-A ribbons (Figure 2D). This trend was similar to that observed for increasing concentrations of VAP-A in membranes, suggesting an effect of VAP-A density on membrane separation in MCS.

### Membrane contact site made with VAP-A can accommodate proteins of different sizes

To assess whether the presence of the bulky ORD (42 kDa) of OSBP modified the MCS architecture, we compared MCS formed with either N-PH-FFAT or full-length OSBP under the same experimental conditions.

Figure 2D and Movies S1-S2 show tomographic reconstruction of an MCS formed by VAP-A and N-PH-FFAT as determined by cryo-ET. The 3D reconstruction revealed a long contact region of ca. 500 nm between a large VAP-A vesicle and a tube, as well as a small contact region of less than 50 nm with a spherical vesicle (see also Figure 3A, 3C and Figures S3 for other examples). As observed in 2D images, the separation between facing membranes was larger when a flattened VAP-A-containing vesicle was in contact with tubes, compared to a spherical VAP-A vesicle; the intermembrane distances were 20-25 nm and 15 nm, respectively. We also found VAP-A ribbons forming contact regions; these were more easily identified in 3D reconstructions than in 2D images. Ribbons, which represent the membrane system with the highest VAP-A density, formed MCS with tubes with an intermembrane distance of 25-30 nm. Both flattened vesicles and ribbons showed an even separation along the whole MCS suggesting homogenous distribution of tethers.

**Figure 3.**
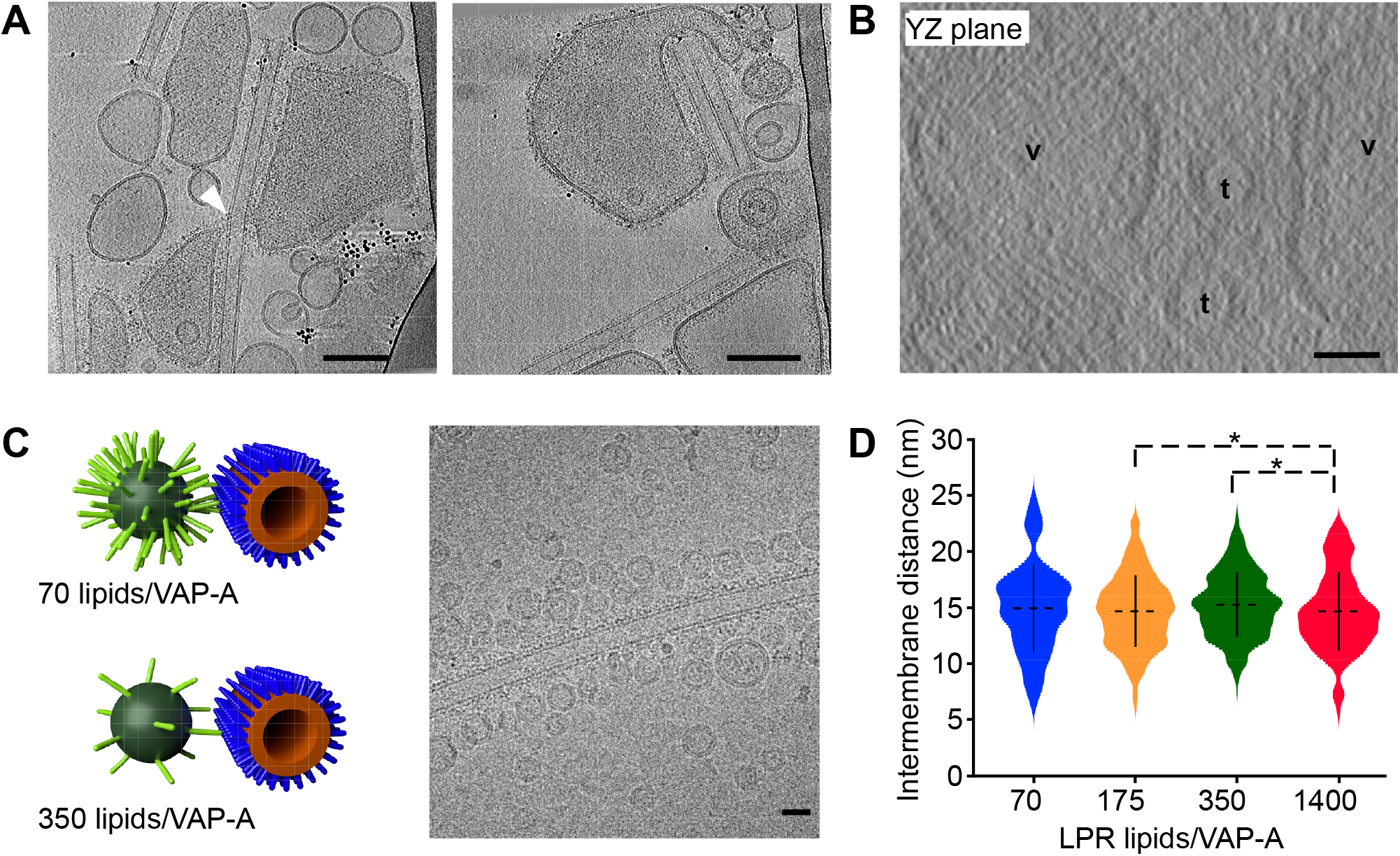
Membrane deformation upon MCS formation. (A) Tomographic XY slice from two tilt series showing that VAP-A vesicles spread along the long axis of tubes, form tongue (left figure, white arrow) around the tubes, but in general avoid the high curvature short axis of tubes (right figure). Bars = 100 nm. (Figure S3A, S3B, Movie S4). (B) Tomographic YZ slice from a third tilt series showing the flat membrane of vesicles (v) facing tubes (t). Bars = 25 nm. (Figure S3C). (C) Small and non-deformable VAP-A proteoliposomes at varying LPR and bound to N-PH-FFAT tubes. Representative MCS with VAP-A vesicles of 30 nm diameter at 70 and 350 lipids/VAP-A mol/mol. Cryo-EM image of small liposomes of VAP-A at LPR 350 lipids/VAP-A forming contact with N-PH-FFAT coated tubes. Bar = 25 nm. (D) Violin plots showing the distribution of distances between small vesicle of VAP-A at varying LPR and N-PH-FFAT tube. The plots show the complete distribution of values analyzed. The horizontal and vertical lines show the average value and the standard deviation, respectively. * indicates p < 0.05 by unpaired t test. A number of 52, 240, 491 and 152 vesicles were analyzed for LPR 70, 175, 350 and 1400 lipid/VAP-A mol/mol, respectively. (Figure S3D).

MCS made with OSBP were more heterogeneous than those observed with N-PH-FFAT. First, the surfaces in contact were smaller even when large VAP-A vesicles were spread along tubes (Figure 2E, Movie S3). Second, membrane separations were shorter than with N-PH-FFAT. In flat contacting regions, where proteoliposomes were aligned along the tubes, we measured intermembrane distances of 10-20 nm, as compared to 15-25 nm for N-PH-FFAT. In line with what we observed for the contacts made at lower VAP-A concentration, this observation suggests that the ORD of OSBP imposes space between proteins, thereby preventing high packing and elongation of VAP-A proteins.

### Membrane remodeling during formation of membrane contact sites

After contact formation between VAP-A vesicles and PI4P-tubes in the presence of OSBP or N-PH-FFAT, we observed significant remodeling of the vesicles, notably their flattening along the major axis of the tubes. When several tubes were involved in the contacts, vesicles were flattened to facing the tubes (Figure 2B, Figures S3 A, B).

In 3D reconstructions of cryo-tomograms, we never observed VAP-A liposomes engulfing a tube. Vesicles were either in reduced contact or formed tongues that partially surrounded the tube (Figure 3A, right, white arrow, movie S4). In some cases, contact-forming vesicles bypassed high curvature areas of tubes (Figure 3A, left). In the YZ plane, the VAP-A membranes facing the tubes were flat and did not bend around the tube, whereas they could clearly curve outside the contact area (Figure 3B). In the presence of two aligned tubes, the vesicle membrane showed a more extended contact but was still flat (Figure S3C). Moreover, VAP-A membranes in contact with the tubes did not show peaks of local deformation, in contrast to what has been reported for tricalbins by *in situ* cryo-EM of ER-plasma membrane contacts in yeast cells (Collado et al., 2019).

We hypothesized that VAP-A molecules concentrated in flat regions of MCS to avoid highly curved membranes. We thus analyzed how MCS formed when high curvature VAP-A vesicles were too tense to be remodeled (Figure 3C). We reconstituted VAP-A at LPR 70 mol/mol and at 4°C instead of 20°C, leading to small and tense proteoliposomes that were homogeneous in diameter (25 ± 10 nm; n=300, 30 images). After incubation with N-PH-FFAT tubes, VAP-A vesicles showed no deformation upon binding along the tubes. Thus, VAP-A can form MCS even between highly curved membranes. However, the distance between the facing membranes was quite short, 15±2 nm (n=52, 4 images). Considering a lipid with a molecular surface of 65Å^2^, a vesicle with a diameter of 20 or 30 nm would contain 3800 or 8700 lipids, respectively, and therefore 52 to 114 VAP-A molecules at the given LPR. When the proteoliposomes were prepared at a high LPR (1400 lipids/VAP-A), leading to on average 2-4 VAP-A molecules per vesicles, the intermembrane distance was similar (ca. 15 nm). These observations suggest that, in an MCS between highly curved membranes, only a few homodimers of VAP-A were involved in the contact (as schematized in Figure 3C). When the VAP-A-containing membrane was deformable, in contrast, proteins seemed to concentrate in flat region of MCS, leading to larger separation between facing membranes, owing to the flexibility of VAP-A.

### 3D models of membrane VAP-A/N-PH-FFAT contact site

To determine the architecture of the contact site at molecular scale, we performed subtomogram averaging. The principle consists in extracting sub-volumes containing identical proteins and associated membranes, followed by averaging them to increase the signal-to-noise ratio. We focused on MCS formed between VAP-A vesicles reconstituted at LPR 70 mol/mol and N-PH-FFAT-decorated PI4P-tubes because they were less heterogeneous than those formed in the presence of OSBP. Protein densities were visible in the contact areas between N-PH-FFAT-containing tubes and hemispherical or flattened VAP-A vesicles (Figure 4A). In some cases, we observed continuous densities joining the two membranes. Out of the contact region, N-PH-FFAT was also seen along the tube as well as VAP-A oriented in and out of the vesicles. These encouraging observations were mitigated by two difficulties. First, the distance between the facing membranes of MCS varied, even in flat regions, in agreement with 2D analysis, suggesting heterogeneity in the organization of protein complexes within MCS. Second, we could not unambiguously attribute the densities to defined protein domains, probably due to the small size of VAP-A and N-PH-FFAT as compared to larger proteins like tricalbins or synatoptotagmins (Collado et al., 2019; Fernández-Busnadiego et al., 2015; Hoffmann et al., 2019).

**Figure 4.**
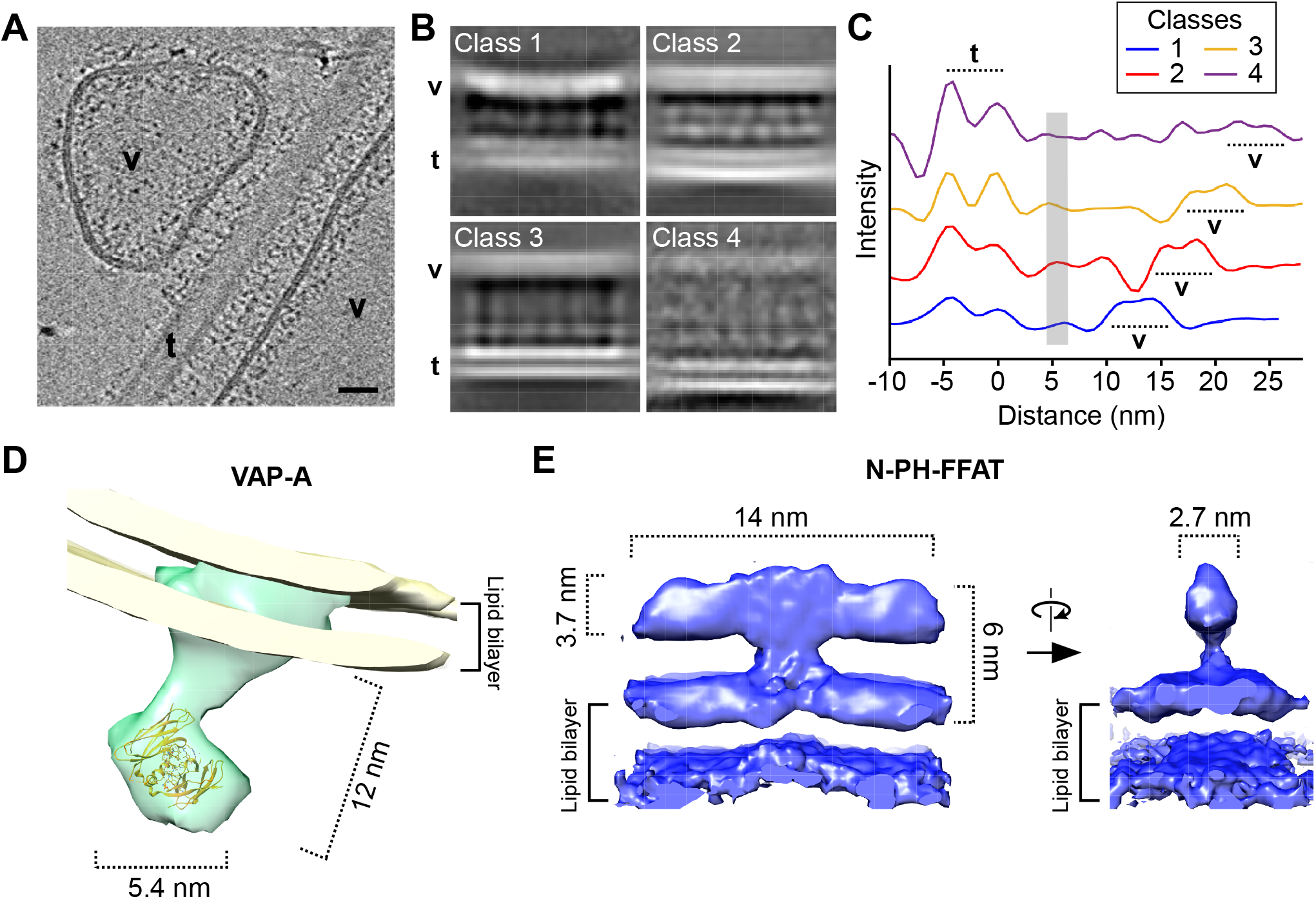
3D molecular architecture of VAP-A N-PH-FFAT in reconstituted MCS. (A) Tomographic slice of VAP-A/NH-FFAT membrane contact site. A hemi-spherical vesicles (v) and a large and flat vesicles (v) contact a tube (t) decorated with N-PH-FFAT bound to. The lipid bilayer of both vesicles is visible while the lipid bilayer of the tube is out of plane. Tiny protein densities tether v and t. Bar = 25 nm. (B) Classes of membrane contact sites after 3D classification. Electron densities of lipid bilayer and proteins are depicted in white. The distance between facing membranes increases from class 1 to class 4. Class 1-3 correspond to MCS formed with VAP-A vesicles and class 4 with VAP-A ribbons. (C) 2D plot of electron densities perpendicular to MCS of classes 1 to 4. The outer leaflets of the tubes were aligned at 0 nm. The two lipid leaflets of the tube are well resolved while the VAP-A membrane is less defined and appears at increasing distance from class 1 to class 4. In all classes, a protein density assigned to N-PH-FFAT domain is found at 5-6 nm from the tubes (gray line). (D) 3D volumes of VAP-A and (E) of N-PH-FFAT at 19.6Å and 9.8 Å resolution, respectively. (Figure S4).

To circumvent these difficulties, we determined major 3D classes of MCS as a function of membrane separation (Figure 4B). The different classes showed electron densities at three levels: (i) the tube membrane, which was well resolved, (ii) the vesicle membrane, which was more diffuse, and (iii) proteins between the two membranes. A plot of the electron densities perpendicular to the membrane plane is shown in Figure 4C, in which we aligned the electron densities at the level of the lipid tube. The density of the VAP-A membrane was at increasing intermembrane distances from class 1 to class 4. In contrast, a protein density was found at a constant distance of 5-6 nm from the tube membrane in all four classes. This density could be attributed to N-PH-FFAT since it was also present in the regions of the tubes not engaged in contacts. This observation suggests that the variable distance within the contacts resulted from the ability of VAP-A molecules to adopt conformations of varying length and/or orientation.

We determined the architecture of the proteins present in class 3, which was the most homogeneous class by sub-tomogram averaging. During the analysis, we observed that the position and densities of N-PH-FFAT were more constant than those of VAP-A (Figure S4). Consequently, we treated the two proteins separately and obtained 3D models of VAP-A and NPH-FFAT at 19.6 Å and 9.8 Å, respectively (Table S1).

VAP-A showed a rod like shape of 12 nm long (Figure 4D). The density attributed to the MSP domain could encompass a dimer confirming a dimeric organization of VAP-A in MCS. The N-PH-FFAT 3D model showed a T-shaped organization with a central stem extending 3 nm from the membrane and joining a 14 nm long and 2.7 nm wide rod-shape structure parallel to the membrane (Figure 4C). The axial symmetry was consistent with a dimer, in line with biochemical observations on OSBP and its N-PH-FFAT construct (Jamecna et al., 2019; Ridgway et al., 1992).

Models of a homologue of the PH domain (aa 80-190 in OSBP) and of the complex between the disordered region G346-S379 containing the FFAT motif (aa 358-361) and VAP-A MSP are available (Lenoir et al., 2010; Furuita et al., 2010). In addition, we have shown that the N-terminal region of OSBP (aa1-80) is disordered (Jamecna et al., 2019). However, there is no structural information on the region connecting the PH domain and the FFAT motif. We previously proposed that this region might form a 10-nm-long rod perpendicular to the membrane composed of two predicted coiled-coils of 30 and 50 aa, that would run parallel to each other. However, such a structure was incompatible with the T-shape observed by cryo EM.

We performed structure predictions with several programs, which suggested similar secondary structures: two helices encompassing T204-S244 (H1) and E250-G324 (H2), followed by a long disordered region encompassing the FFAT motif (aa 325-408) (Figure 5A, S5B). We hypothesized that these two helices made the main contribution to the electron density of the T structure, while the PH domains would lie on the membrane, and the unstructured parts were not visible at this resolution. To test this model, we expressed and purified an OSBP construct encompassing aa 199-324, hereafter termed OSBP Central Core (CC). In agreement, the circular dichroism spectrum of this construct showed 86% of alpha-helical content, confirming the presence of two helices, whereas gel filtration chromatography revealed a dimeric organization (Figure 5B, 5C).

**Figure 5.**
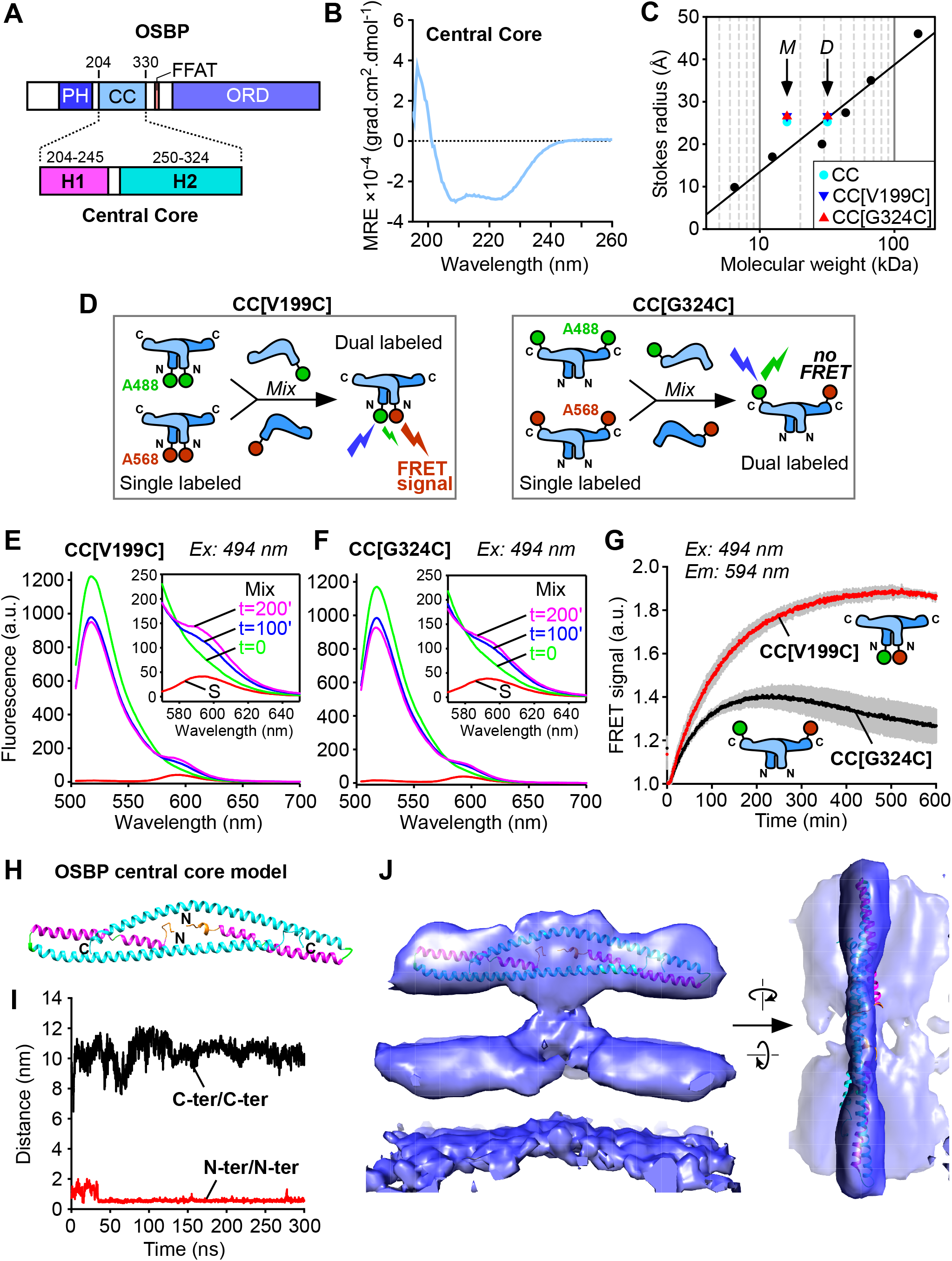
3D architecture of the central core domain of OSBP. A) Domain organization of OSBP with its central core (CC) region and the predicted H1 and H2 helices. (Figure S5) (B) Circular dichroism spectroscopy of OSBP CC region. CD spectra minima at 208 and 222 nm indicative of a significant α-helical content. (C) Stokes radius versus MW of OSBP CC constructs (WT, light blue dot; V199C/C224A/C276A, blue triangle; C224A/C276A/G324C, red triangle) and standards (black dots) as determined by gel filtration. The arrows indicate the expected positions for monomers (M) or dimers (D) according to their MW. (D) Principle of OSBP CC monomer exchange experiment as analyzed by FRET (E-F) spectra and (G) kinetics analyses of monomer exchange between AF-488 and AF-568 labelled OSBP CC region. Labelling was performed an a Cys introduced either on the N-ter (V199C) or the C-ter (G324C) of a construct having endogenous Cys substituted to Ala (C224A/C276A). (H) Structure of the dimer OSBP CC after 300ns MD simulation colored by chains. (I) Plot of the distances between C-termini and N-termini along the MD trajectory (300ns). (J) Fit of the dimer OSBP CC after 300ns MD simulation in the 3D EM model of N-PH-FFAT.

In order to better understand the organization of the CC region of OSBP, we designed an experiment to probe intermolecular fluorescence transfer (FRET) within dimers (Figure 5D). We mixed a dimer population where each chain was labelled with a donor probe with a dimer population where each chain was labelled with an acceptor probe. The exchange of monomers over time should lead to the formation of hybrid dimers harboring monomers of different colors. If the aa chosen for fluorescent labelling on one chain was close to the cognate aa on the other chain, this should result in a FRET signal. This strategy allowed us to the test the T shape model: the two N-ter should be close together, whereas the two C-ter should be remote. In contrast, if this region adopted a parallel coiled-coil structure, all aa pairs should be close together along the structure and a FRET signal should be observed whatever the position chosen for labeling.

We replaced the endogenous cysteines of OSBP CC (C224 and C276) by alanines and we introduced a cysteine either at the N-terminus (CC[V199C] construct) or at the C-terminus (CC[G324C] construct). All mutants behaved similarly by gel-filtration, suggesting no effect on the structure (Figure 5B). After purification, each construct was labelled with a donor probe (AF-488) or an acceptor probe (AF-568). When the CC[V199C] labeled dimers were mixed, a FRET signal was observed, indicating proximity of the N-termini in the dimer. Monomer exchange occurred with a time course of several hours (half-time = 100 min) indicating a very stable dimeric structure. In striking contrast, a much lower FRET signal was observed when mixing the CC[G324] labeled dimers, suggesting that the C termini were not close within the dimer (Figure 5E-G).

We generated 3D models of 195-335-OSBP with Robetta (Song et al., 2013). All models shared similar organization with H1 and H2 helices, H2 helix being continuous (e;g. model 1) or discontinuous (e.g. model 2) (Figure S5B). We used Molecular Dynamics (MD) simulations to analyze the dimer stability and found that model 1 formed a stable dimer (Figure 5H). The distances between C-termini and N-termini along the MD trajectory (300ns) are depicted Figure 5I. Within the dimer, the N-termini and the C-termini were close and distant, respectively, in agreement with the FRET experiments. The long H2 helix that contains the large dimerization region (261-296) proposed by (Ridgway et al., 1992) participates to the formation of the dimer. The 14 nm long dimer was fitted in the horizontal part of the T of N-PH-FFAT, the stem of the T remaining to be assigned.

## DISCUSSION

We have reconstituted a contact site between two model membranes and a protein complex made by the general ER receptor, VAP-A, and the founding member of the ORP family, OSBP, or its N-PH-FFAT construct. Cryo-EM and cryo-ET analysis show MCS that are separated by distances ranging from 10 to 30 nm. After 3D classification of the different MCS sub-regions, we obtained a 3D model of each protein within the MCS. We suggest that VAP-A is a highly flexible protein, allowing MCS formation of varying intermembrane distance, which increases with protein density. N-PH-FFAT appears more rigid owing to a central, dimeric, and helical T-shape region. This T geometry should keep the two ORD in the OSBP dimer well separated and might be favorable for their movements between the two facing membranes during lipid exchange.

### VAP-A is a flexible protein and adapt to its interactors within the MCS

When VAP-A is not engaged in a contact, it appears as a flexible protein, extending its MSP domain at increasing distances that correlate with protein density. At a medium density, VAP-A shows various orientations with regards to the membrane plane, with its MSP domain exploring a 10 nm-thick region. At a very high density that leads to ribbons-like membranes, the MSP extends to 17 nm away from the membrane (Figure 1F). Structure predictions propose that two unstructured regions flank the predicted coiled-coil domain of VAP-A: region 135-171 between the MSP domain and the coiled-coil, and region 208-226 between the coiled-coil and the TM domain (Figure S5A). These regions, which are also present in VAP-B, might provide VAP-A/B dimers with bending and stretching capacity, allowing exploration of a larger space than a rigid molecule on the surface of the membrane (Figure S5C).

A most striking feature of VAP-A/B is their ability to interact with a myriad (currently estimated at ≈ 100) of cytosolic protein domains. VAP-A/B are in fact the only known receptors that anchor proteins on the ER surface. Importantly, the proteins that are recruited by VAP-A/B have very different domain organization and function. Some effectors are simply retained at the ER, others bridge the ER to various organelle membranes, as is the case for OSBP. For example, the Opi1 repressor interacts with the yeast VAP-A homologue as well as phosphatidic acid at the ER surface, whereas MIGA2 bridges ER to mitochondria (Freyre et al., 2019; Gaspar et al., 2017). These different molecular contexts might require different orientations and positioning of the MSP domain of VAP-A/B. Therefore, we propose that the structural flexibility of VAP-A/B enables a large spectrum of protein assemblies. In addition, fast rotation of the MSP domain might favor rapid association kinetics, akin to what has been found for golgins, where flexible regions between coiled-coil regions have been shown to be essential for the capture of vesicles on the surface of the Golgi (Cheung et al., 2015). In Ist2, a yeast MCS tether, a long intrinsically disordered region is also responsible for membrane tethering, spatial freedom and protein recruitment (D’Ambrosio et al., 2020; Kralt et al., 2015). Model studies of the interaction between streptavidin-coated vesicles and flat surfaces coated with either flexible PEG-biotin or rigid casein-biotin ligands show that molecular flexibility can increase the rate of adhesion between two surfaces by three orders of magnitude (Cuvelier and Nassoy, 2004).

When VAP-A is engaged in a MCS with N-PH-FFAT, 3D classification reveals that, again, VAP-A is variable in length and determines the separation between membranes; in contrast, N-PH-FFAT displays a fixed length (Figure 4C). The intermembrane distance increases with the concentration of VAP-A in MCS from 10 nm when a few VAP-A molecules are reconstituted in small vesicles to 30 nm when VAP-A molecules are densely compacted in ribbons (Figure 2C, Figure 3D).

VAP-A forms MCS with OSBP or its membrane tethering moiety N-PH-FFAT. However, less VAP-A molecules are engaged within MCS formed with OSBP than N-PH-FFAT as shown by the shorter intermembrane distance suggesting that the bulky ORDs decrease the accessibility of MCS to VAP-A (Figure 2D). VAP-A is thus strikingly different from a rigid tethering protein that imposes a fixed distance between the facing membranes. In that case, as shown by model mixtures of binding and non-binding proteins of variable lengths, a difference in length of only 5 nm of the non-binding proteins is sufficient to drive exclusion from MCS (Schmid et al., 2016). The much larger range of intermembrane distance in VAP-A mediated MCS suggests that VAP-A adapts its density and dimension to the size of its interactor within a contact area (Figure S5D).

At ER-Golgi MCS, VAP-A recognizes several lipid transfer proteins including OSBP, CERT, Nir2 and FAPP2, which deliver cholesterol, ceramide, PI and glucosylceramide to TGN respectively (Mesmin et al., 2019). They all encompassed a PH or a LNS2 (Nir2) domain that binds PI4P at the Golgi and a FFAT motif. As a result, these LTPs could bind simultaneously to both organelle membranes and thus act as bridging factors. It is not known if VAP-A has different affinity for these different LTPs and if all LTP could coexist within the same MCS. However, our results suggest that the flexibility of VAP-A could support the coexistence of different partners involved in lipid transfer within the same MCS. Similarly, it has been shown that the phosphoinositide transfer protein Nir2 is recruited at ER-PM neuronal contact sites made by VAP-A and the potassium channel, Nir2 and Kv2:1 bearing a FFAT and FFAT-like motif, respectively. Such tripartite ER-PM junction is proposed to play a role in the phophoinositide homeostasis in brain neurons (Kirmiz et al., 2019).

### VAP-A proteins concentrate in flat membrane regions of MCS

During the formation of the MCS, large VAP-A vesicles spread along the major axis of the PI4P containing tubes until they were no longer deformable (Figure 3A). This suggests the participation of several VAP-A/N-PH-FFAT tethering complexes in an MCS, which could compensate for the moderate affinity (Kd 2 μM) of FFAT-bearing proteins for MSP (Furuita et al., 2010). On the other hand, we observed no wrapping of VAP-A vesicles around the highly curved tubes, which would require strong interaction between VAP-A and N-PH-FFAT or OSBP. When membrane curvature is too high, like in small and tensed proteoliposomes, the separation is small and independent of the initial density of VAP-A in the proteoliposomes (Figure 3C). Thus, the high curvature in small proteoliposomes prevents VAP-A molecules from forming a dense protein area as observed in flat membranes (Figure S5D). In situ cryo-EM has shown that Scs2, the yeast ortholog of VAP-A, also accumulates in ER sheets, while tricalbins accumulate in tubular ER (Collado and Fernández-Busnadiego, 2017; Hoffmann et al., 2019). This difference was attributed to the cylindrical shape of Scs2 with a single transmembrane helix and an elongated cytosolic domain as compared to the conical shape of tricalbins, which insert in the membrane *via* a hairpin and display bulky C2 domains.

Once in flat regions, how VAP-A concentrate in MCS remains an open question. Clusters of VAP-A have been reported as a consequence of domains of Kv2.1 potassium channel in the PM and binding of VAPs to form ER-PM contact site (Johnson et al., 2018). Concentration of VAP-A might also result from geometrical constraints: when an elementary VAP-A/N-PH-FFAT pair forms, similar pairs should concentrate in the vicinity as the intermembrane distance is optimal for assembly. VAP-A might also concentrate in lipid nano-domains present in ER as reported recently for other MCS (King et al., 2019). However, our liposome composition was not prone to form membrane domains. Whatever the process of concentration of VAP-A, our results suggest that it should have an impact on the separation of facing membranes within MCS.

### The central core of OSBP forms a T shape

The T-shaped organization of tethering region of OSBP was unanticipated. We previously suggested that the overall architecture of the region between the PH domain and FFAT motif (aa ≈ 200-325) forms a ≈ 10 nm long rod perpendicular to the membrane plane because it contains two predicted parallel coiled-coils. Instead, cryo EM analysis shows a 14 nm elongated domain parallel to the membrane. A short ≈ 3 nm stem, which probably follows the two PH domains, connects this elongated structure to the membrane. Biochemical analysis indicates that the region between the PH domain and FFAT motif is dimeric, alpha-helical and has its N-termini very close together, while its C-termini are far away (Figure 5). The separation between the two C-termini is rather in favor of an antiparallel organization. Regardless the exact molecular structure, which will require further studies, the consequence of this organization is that the tethering and lipid transfer moieties of each monomer are separated by a large distance, akin to the jib of a tower crane. Moreover, the two ORDs of the OSBP dimer should be well separated from each other, which might facilitate their independent movements.

Following the T domain, a long sequence G324-R408 is predicted to be unstructured (Figure S5B). The first part of this sequence, G324-N357, i.e. 31 aa before the FFAT motif, might extend up to 10 nm beyond the T domain to interact with VAP-A (considering a 3.5 Å step per residue). As a result, the orientation and position of the T domain relative to VAP-A might vary. This is what we observed when analyzing a 3D class defined by the fixed intermembrane distance: the VAP-N-PH-FFAT complexes were not at fixed positions thus limiting the resolution of the whole complex. The second part of the sequence A362-R408 (= 47 aa), which unifies the FFAT motif to the ORD, could further extend 14 nm away and should define the action range of the ORD between the MSP of VAP-A and the two facing membranes. This ball-in-chain geometry could allow a movement of up to 28 nm of the ORD between the ER and TGN membranes, which might be enough to exchange lipids between the two facing membranes, considering that MCS at the ER TGN interface have an intermembrane distance of 5-20 nm (Venditti et al., 2019). In addition, the flexibility of VAP-A as revealed here by cryo EM could further facilitate the movement of the ORD.

We have previously shown that the intrinsically disordered N-terminal region of OSBP controls its density under confined conditions, which facilitates lateral diffusion of proteins within MCS (Jamecna et al., 2019). Here, we highlighted the flexible properties of VAP-A and the ordered and disordered domains of OSBP. Thus, the molecular actors of VAP-A/OSBP MCS carry interfacial recognition information, and also contain regions that are intrinsically disordered and flexible, which is likely crucial for MCS dynamics and supramolecular assembly.

## ACKNOWLEGMENTS

We are grateful to A. Bertin for 2D image analysis, P. Cuniasse and P. Sens for fruitful discussions, D. Woolfson for feedback on coiled-coil analysis, N. Charmel for the picture of Figures 2A and 3C and L. Duchesne for providing PEG-coated gold beads. We thank G. Schoehn and the EM facilities at the Grenoble Instruct-ERIC Center (ISBG; UMS 3518 CNRS CEA-UGA-EMBL) with support from the French Infrastructure for Integrated Structural Biology (FRISBI; ANR-10-INSB-05-02). We are deeply grateful to W. Hagen for the quality of the cryo-electron tomography data acquired at the cryo-electron microscopy platform of the European Molecular Biology Laboratory (EMBL) in Heidelberg. We thank the Cell and Tissue Imaging core facility (PICT IBiSA), Institut Curie, member of the French National Research Infrastructure France-BioImaging (ANR10-INBS-04). This work was supported by CNRS, Institut Curie, and the Agence Nationale de la Recherche (ANR-15-CE11-0027-02). This work was also supported by iNEXT, project number iNEXT PID7315, funded by the Horizon 2020 program of the European Union. E. de la Mora was funded by a grant from ANR (ANR-15-CE11-0027-02) and by a grant from the Labex CellTisPhysBio (ANR-11-LABX-0038, ANR-10-IDEX-0001-02). The Antonny lab is supported by a grant from the Fondation pour la Recherche Médicale (Convention DEQ20180339156 Equipes FRM 2018), by the LABEX signalife (ANR-11-LABX-0028-01) and by the university Côte d’Azur (IDEX académie 4 – masters environnés).

## AUTHOR CONTRIBUTIONS

Conceptualization, MD, BM, BA and DL; Purification of LTPs and lipid transfer assays JB, JP, BM and of VAP-A, MD and JM. Cryo-EM, cryo-ET, ADC, MD and DL. Biochemical analysis JB, BM, BA and MD. Image analysis EM, DCD. Modelisation DL, RG. Writing – Original Draft, DL and MD; Writing – Review & Editing, DL and BA. Funding Acquisition, BM, BA and DL.

---

**TABLE S1.**
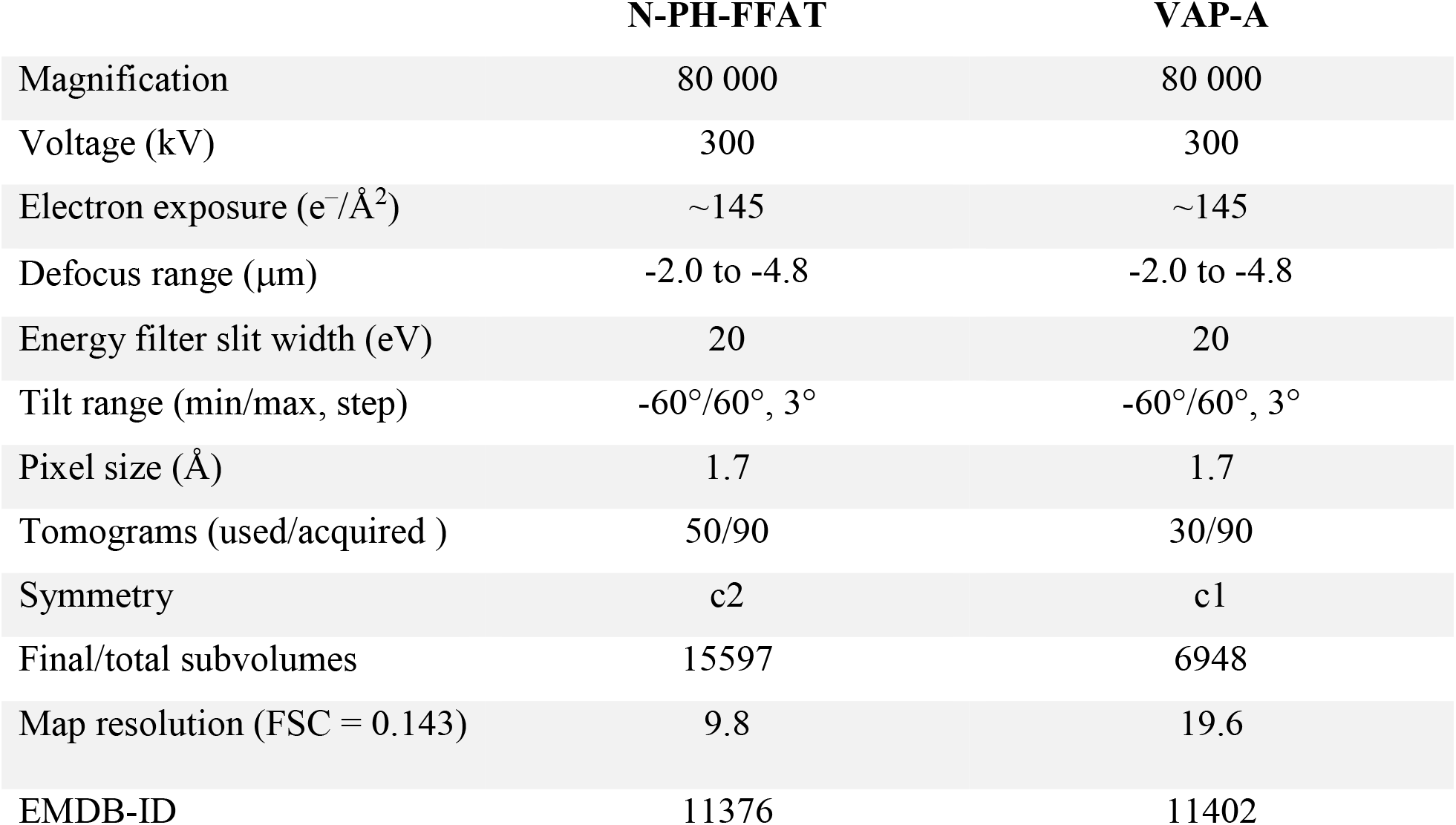
Data collection and image processing table

## STAR Methods

### KEY RESOURCES TABLE

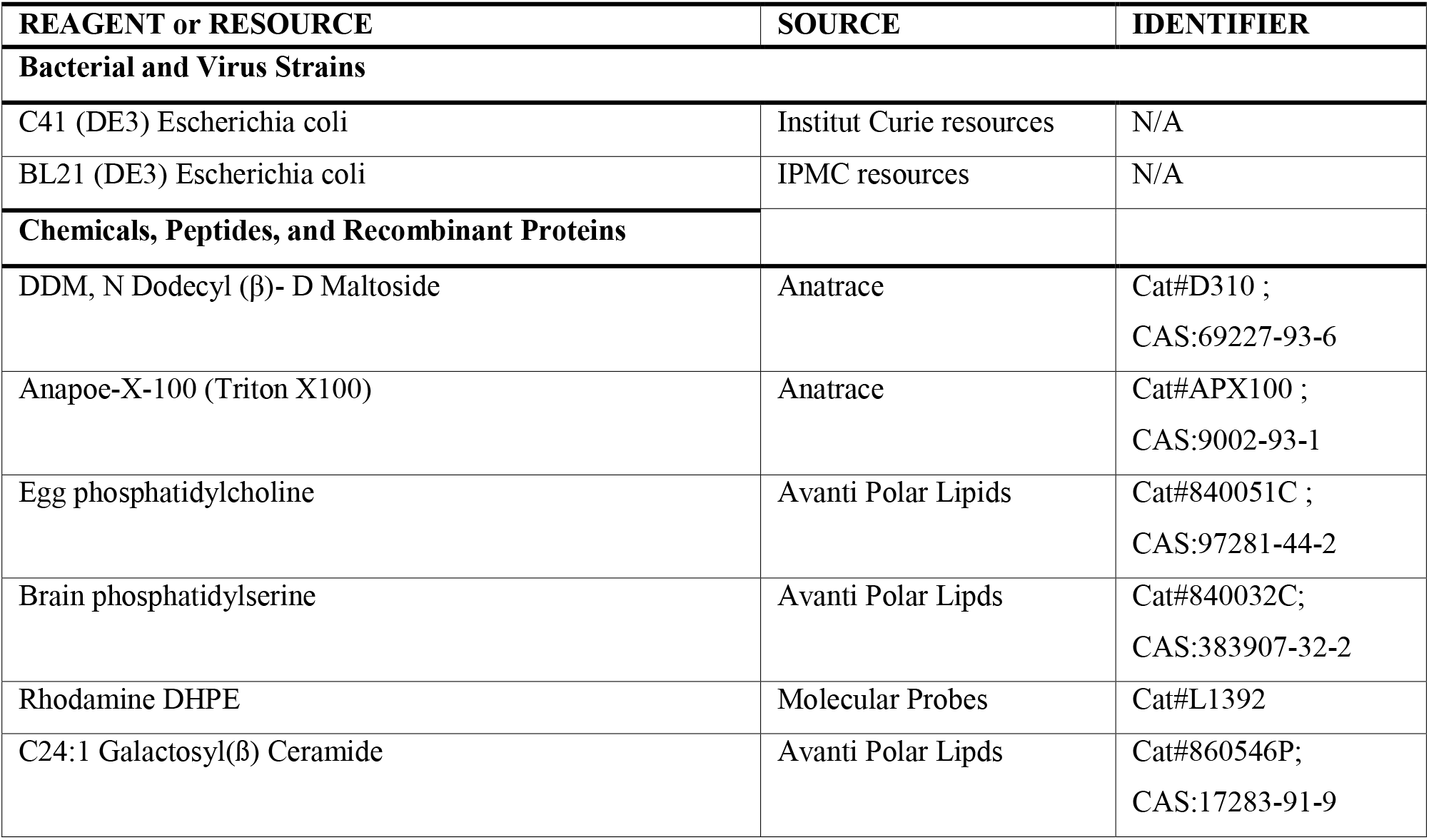
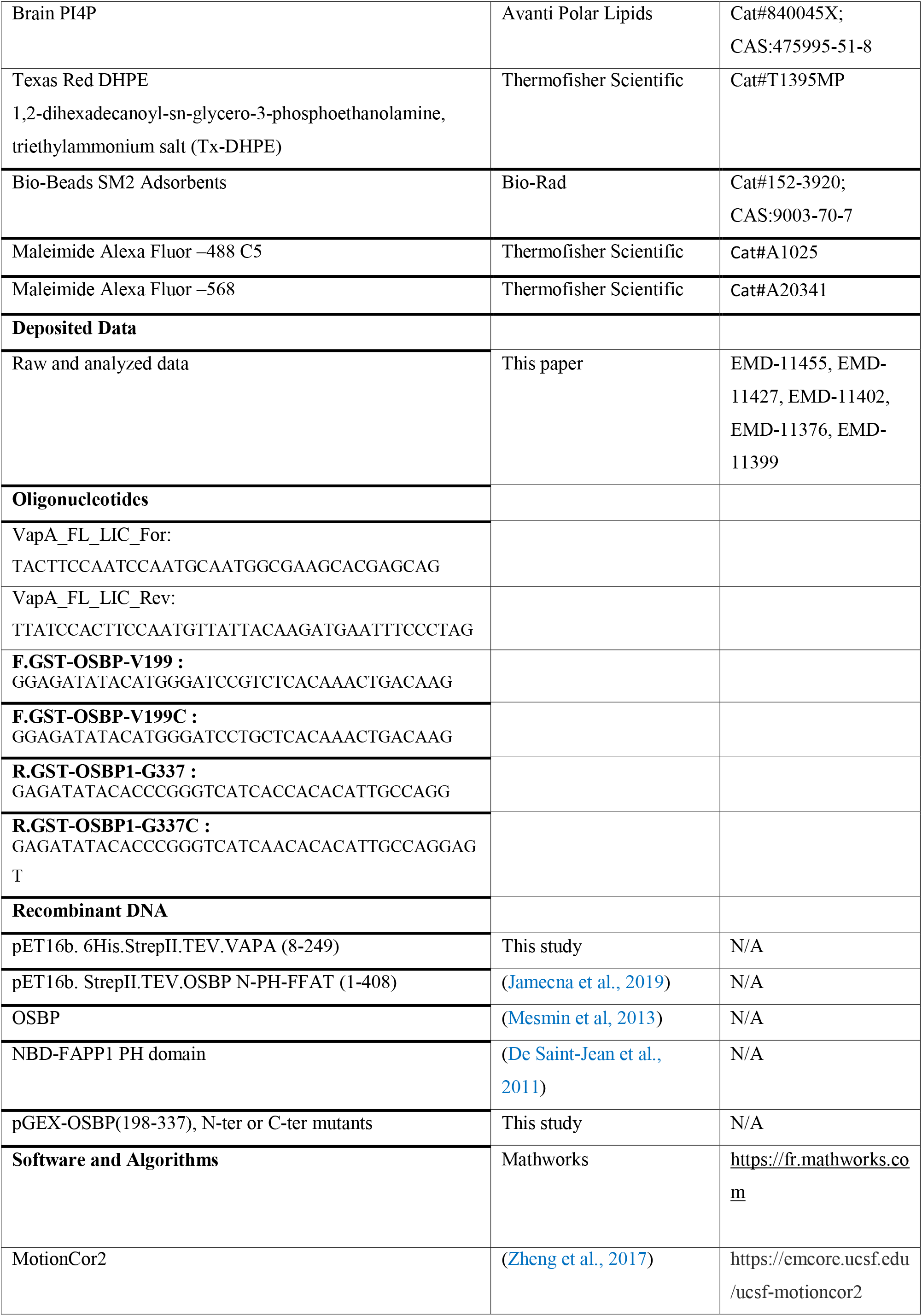
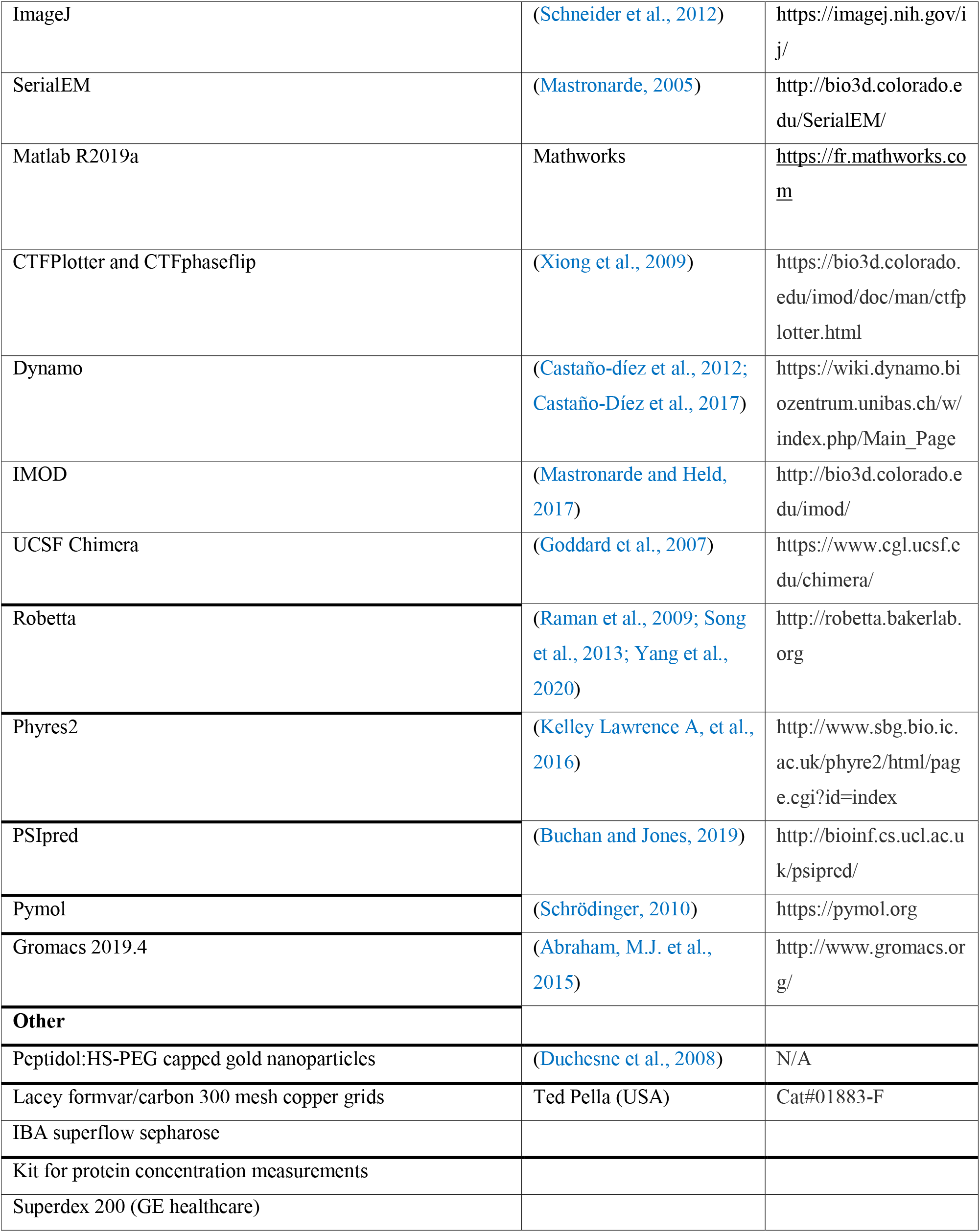

### RESOURCE AVAILABILITY

Lead Contact: Further information and requests for resources and reagents should be directed to and will be fulfilled by the lead contact Daniel Lévy).

#### Materials Availability

Plasmids generated in this study are available upon request to the lead contact.

#### Data and Code Availability

The data supporting the findings of this study are available within the article and its supplementary materials. Tomograms and generated maps were deposited to EMDB (https://www.ebi.ac.uk/pdbe/emdb/) with accession numbers EMD-11455, EMD-11438, EMD-11402, EMD-11427, EMD-11399, EMD-11376.

### EXPERIMENTAL MODEL AND SUBJECT DETAILS

#### Microbe Strains

Recombinant Vap-A proteins were expressed in C41 (DE3) *E. coli* strain, which were grown in 2xYT medium at 30 °C. Recombinant OSBP proteins were expressed in Bl21 (DE3) *E. coli* strain, which were grown in LB/Ampicilin medium at 37 °C and induced at 16°C.

### METHOD DETAILS

#### Reagents

Egg PC, brain PS, brain PI(4)P, C24:1 Galactosyl(ß) Ceramide, Texas Red 1,2-dihexadecanoyl-sn-glycero-3-phosphoethanolamine, triethylammonium salt (Tx-DHPE) were from Avanti Polar Lipids. Anapoe-X-100 (Triton X100), N-Dodecyl-β-D-maltoside (DDM) was purchased from Anatrace. Peptidol:HS-PEG capped gold nanoparticles of 6 nm were prepared by Dr. L. Duchenes (Duchesne et al., 2008); 50 mesh BioBeads were purchased from Bio-Rad and prepared according to (Rigaud et al., 1997). Lacey carbon electron microscopy grids were purchased from Ted Pella (USA).

#### Plasmids and proteins expression

Human VAP-A (8-249) fragment was cloned into pET-His6-TEV-LIC then sub-cloned into pET16b for inducible expression as N-terminal His-StrepII-TEV-Vap-A. Vap-A expression was performed in C41 (DE3) *E. coli* strain. Full length VAP-A expression was performed in presence of 0.5 mM IPTG for 4 hours at 30 °C.

OSBP central region (198-337) were PCR amplified as either (V199C/C224A/C276A) or (C224A/C276A/G337C) mutants and sub-cloned via BamH1/Xma1 sites into pGEX-4T2 for expression as GST-tag proteins. Proteins were expressed in Bl21gold *E. coli* after induction with 1mM IPTG and overnight culture at 16°C.

#### Proteins purification

1. VAP-A: VAP-A expressing cells were collected and passed twice through a Cell Disrupter at 2k bar pressure. The resulting cell lysate was centrifuged to purify membranes. Membranes were solubilized for 2h at 4°C with DDM:protein ratio of 2.5 (w/w) and then centrifuged to collect solubilized proteins. VAP-A was purified on Strep Tactin beads (IBA superflow sepharose) in batch. Eluted proteins were incubated overnight with TEV protease to eliminate the double tag, followed by purification on a Superdex 200 (GE healthcare) size exclusion chromatography in Tris 50 mM pH 7.5, NaCl 150 mM, DDM 0.03mol/, 10% glycerol. Purified VAP-A was collected at concentration ranging from 1.5 to 2 mg/ml without any additional concentration step, flash frozen in liquid nitrogen and stored at −80°C.
2. OSBP and N-PH-FFAT were purified according to (Jamecna et al., 2019; Mesmin et al., 2013). GST-OSBP (199-324) mutants expressing cells were pelleted and resuspended in 50 mM Tris (pH7.5), 300 mM NaCl, 5 mM DTT buffer. The bacteria were lysed through a Cell Disrupter at 2k bar pressure and cell lysate was ultra-centrifuged. Soluble GST-tagged proteins were first purified on Glutathion-Sepharose 4B (GE healthcare) beads, eluted after thrombin clivage, and then further purified on a MonoS HR5/5 column (GE healthcare). Proteins were eluted from the column with a 0 to 1M NaCl gradient (25 column volume) in 25 mM MES (pH 6.0), 3 mM DTT buffer. Protein fractions were pooled, concentrated on Amicon Ultra 4 (10kDa cut off), injected on a Superpose 12 HR 10/30 column (GE healthcare) and eluted with Tris 25 mM pH 7.5, NaCl 120 mM, DTT 3 mM buffer. Proteins fractions were pooled, concentrated on Amicon Ultra 4 (10kDa cut off), and supplemented with 10% glycerol before flash freezing in liquid nitrogen and stored at −80°C.

#### Analytical Gel filtration

Purified proteins (100 μl, 10 μM) were applied to a Superdex 75™ column (GE Healthcare) and eluted at a flow rate of 0.5 ml.min^-1^ in 25 mM Tris pH 7.5, 120 mM NaCl and 1mM DTT. The column was calibrated using the following standards (MW/Stokes radius): Bovine serum albumin (67 kDa/3.5 nm), Ovalbumin (43.5kDa/2.7 nm), Carbonic anhydrase (25 kDa/2.1 nm), Cytochrome C (12.4 kDa/ 1.7 nm) and Aprotinin (6.5 kDa/0.9nm). The elution volume and Stokes radius of the standards were used to establish a first calibration curve, from which the Stokes radius of the OSBP constructs were determined. Thereafter, we plotted the Stokes radius as a function of MW for both protein standards and for OSBP constructs.

#### Fluorescent labelling

DTT present in OSBP (199-324) CYS mutants was removed by buffer exchange on a NAP-5 column (GE healthcare). Proteins were incubated with a 10-fold mol excess of Alexa dyes (AlexaFluor-488 dye and AlexaFluor-568) for 1 min at room temperature and then 30 min in ice. Reaction was stopped by addition of a 100-fold excess of L-cystein. Excess unbound dye and L-cyst was removed by elution on a NAP-5 column (GE healthcare).

#### Circular Dichroism

CD spectroscopy was performed on a Jasco J-815 spectrometer at room temperature with a quartz cell of 0.05 cm path length. Each spectrum is the average of 10 scans recorded from 195 to 260 nm with a bandwidth of 1 nm, a step size of 0.5 nm and a scan speed of 50 nm min−1. The purified construct OSBP (198-324) was dialyzed in Slide-A-Lyzer dialysis cassette (Thermo) against KCl 150 mM, Tris 10 mM pH 7.5, DTT 1 mM and was used at 48 μM. Control spectra of buffer alone was subtracted from the protein spectra.

#### FRET measurement

Fluorescence Resonance Energy Transfert (FRET) reactions were determined by mixing AlexaFluor-488 labeled to AlexaFluor-568 OSBP constructs (100 nM each) in a stirred cuvette containing 600 μl HKM buffer upon excitation at 494 nm (1 nm bandwith) and spectra emission from 500 to 700 nm (5 nm bandwith). Spectrum were measured every 10 min for V199C-337 construct and every 5 min for 198-N337C (total 20 spectra each). For kinetic measurements, emission was set at 594 nm (5 nm bandwith).

#### Reconstitution of VAP-A in proteoliposomes

VAP-A was reconstituted in 9.5/0.5 w:w EPC/PS lipid mixture by detergent removal using Bio-Beads (see reviews (Rigaud and Lévy, 2003; Rigaud et al., 1998, 1995)). Briefly, DDM solubilized VAP-A was mixed at room temperature with EPC/PS liposomes solubilized in Triton X100, detergent/lipid ratio of 2.5 w:w, in 50 μl volume. Detergent was removed by additions of wet BioBeads at Bio-Beads/detergent ratio of 20 w:w 2 hours/20 w:w 1h/20 w:w 1h at room temperature or at a ratio of 40 w:w for 2 hours and a second addition of 20 w:w for 2 more hours at 4°C. This amount of Bio-Beads allowed a complete detergent removal (Rigaud et al., 1997). Reconstitution buffer was 50 mM Hepes pH 7.4 and 120 mM potassium acetate (HK buffer), 1 mg/ml lipid, 2.5 mgr/ml Triton X100. Protein was added at lipid/protein ratio from 70 to 2800 mol/ mol. After reconstitution, proteoliposomes were kept at 4°C for a maximum of two days before use.

#### VAP-A incorporation into proteoliposome

VAP-A was reconstituted as described above in a lipid mix EPC, bPS 9.5:0.5 w/w and 0.5 w/w of fluorescent Tx-DHPE, at LPR 1400 mol/mol. Then 100 μl of VAP-A proteoliposomes were mixed with freshly prepared ice cold 50% w/w sucrose in gradient buffer (Tris pH 7.4, 50 mM, NaCl 150 mM) to get final 575 μl of 30% sucrose concentration fraction and loaded at the bottom of a gradient tube (2.5 ml, Ref.347357 Beckman Coulter). Then 575 μl fraction of 20, 10 and 5 % sucrose concentration were carefully loaded one after the other to create a discontinuous gradient. Gradient preparation was performed at 4°C to allow a good separation of sucrose fractions and avoid any mixing.

#### PI4P transport

For PI(4)P transfer assay, Golgi-like liposomes with 2 mol % Rho-PE and 4 mol % PI4P (250 μM lipids) were incubated in HKM buffer with 0.3 μM NBD-FAPP1 PH domain (NBD-PH) and with 0.17 μM VAP-A reconstituted at LPR 40 in proteoliposomes (La, 250 μM lipid). PI(4)P transport was followed by measuring the NBD emission signal at 530 nm (bandwidth 10 nm) upon excitation at 460 nm (bandwidth 1nm) in a JASCO fluorimeter. At the indicated time, 0.1 μM OSBP were added.

#### Preparation of galactocerebroside tubes

Galactocerebroside (Galcer) tubes doped with PI4P were prepared according to (Wilson-Kubalek et al., 1998). Briefly, a mixture of Galcer, EPC, brain PS, brain Pi4P was dried under vacuum. Galcer/EPC/PS/Pi4P (80/10/5/5) tubes were formed at room temperature after resuspension of the dried film at 5 mg/ml lipid concentration in HK buffer followed by 5 cycles of 10 minutes vortex, 2 minutes at 40°C. GalCer tubes were aliquoted and stored at −20°C.

#### Formation of in vitro membrane contact sites

VAP-A/N-PH-FFAT and VAP-A/OSBP MCS were formed directly on the cryo-EM grid at room temperature. VAP-A proteoliposomes were diluted at 0.05 mg/ml in HK buffer. Galcer tubes were diluted at 0.15 mg/ml and mixed with NPHFFAT or OSBP at 70 lipids protein mol/mol ratio. A 2 μl drop of NPHFFAT or OSBP tubes was immediately loaded on the grid and let incubate for about 1 min, followed by the addition of 2 μl VAP-A liposomes. After 30 seconds incubation, grid was blotted and immediately frozen. For cryo-tomography experiments 6 nm gold beads were added before plunging. This protocol was used for all reconstitutions of VAP-A with LPR ranging from 70 to 2800 mol/mol.

#### Cryo-EM experiments

Lacey carbon 300 mesh grids (Ted Pella, USA) were used in all cryo-EM experiments. We found that lacey networks with holes ranging from 20 nm to few microns entrapped different reconstituted material more than calibrated Quantifoil grids. In all experiments, blotting was carried out on the opposite side from the liquid drop and plunge frozen in liquid ethane (EMGP, Leica, Germany). Samples were imaged using different electron microscopes. Data and 2D images depicted in Figure 1C, Figures S1A-L, Figures S2A-E, Figure 3C and Figure S3D were acquired with a Tecnai G2 (Thermofisher, USA) Lab6 microscope operated at 200 kV and equipped with a 4k x 4k CMOS camera (F416, TVIPS). Image acquisition was performed under low dose conditions of 10 e^-^/Å^2^ at a magnification of 50,000 or 29,500 with a pixel size of 2.1 or 3. 6 Å, respectively. Plots depicted in Figure 1E, Figure 3D were derived from images taken with the Tecnai G2 electron microscope. Data and 2D images depicted in Figure 1E-F, Figure 2B, plot Figure 2B were acquired with a Polara (Thermofisher, USA) FEG 300 kV microscope with a K2 Gatan camera in counting mode with 40 frames during 6 sec, total dose 80 e^-^ and at 0.96 Å /px.

#### Cryo-ET data acquisition

Tilt series from samples of VAP-A/OSBP and VAP-A/N-PH-FFAT were acquired with a Titan Krios microscope (Thermofisher, USA) operated at 300 keV and equipped with a Quantum post-column energy filter and a Gatan K2 Summit direct detector. Tilt series, consisting on 41 images, were collected under the dose-symmetric scheme (Hagen et al., 2017) in the angular range of ±60°, with angular increment of 3°, defocus range between −2.0 and −4.8 um and pixel size of 1.7 Å. Every image was composed of 10 frames with a total dose of 3.5 e^-^/ Å^2^. The total dose for each tilt series was 143.5 e^-^/Å^2^. High resolution movies were aligned with MotionCor2 (Zheng et al., 2017), and the average defocus was estimated with CTFPlotter and corrected with CTFphaseflip (Xiong et al., 2009). The resulting micrographs were dose-filtered according to (Grant and Grigorieff, 2015) by means of Matlab scripts (https://github.com/C-CINA/TomographyTools).

During optimization of cryo-ET samples, we observed that protein A gold beads bound with high affinity to VAP-A. To reduce binding, PEG-coated gold beads were added to samples employed for cryo-ET (Duchesne et al., 2008). We observed that much more gold beads were at the proximity of MCS formed by OSBP than with N-PH-FFAT. Cryo-ET (see Figure 3D) revealed that less molecules of VAP-A were engaged in contacts with OSBP and thus more accessible for binding gold particles. However, this prevents further analysis by sub-tomogram averaging of OSBP-VAP-A MCS. Data collection are presented Table S1.

#### 2D image analysis

In 2D cryo-EM images, the lipid bilayer of the vesicles or the tubes are identifiable and serve as landmarks for distance measurements. In the plot Figure 1E, the lengths of the extramembraneous domains of VAP-A are reported from the electron densities extending at the largest distances from the membrane. A number of 50 vesicles (n=78 measurements), 45 ribbons (n= 421 measurements) and 6 onions (n=78 measurements) have been analyzed. In the case of onions, we consider VAP-A length as half of the intermembrane distance although we have no details on the molecular organization of VAP-A homotethers. In Figure 2B contact areas are identified by the presence of deformed and spread vesicles in contact with tubes and electronic densities between membranes. In the case of spherical vesicles with low density of VAP-A, vesicle involved in contacts are identified by their proximity, < 30 nm, to tubes. In addition, to avoid selecting vesicles close to tubes due to too high a concentration of material in the holes, only areas where tubes and vesicles are sparsely distributed in the holes of the lacey are considered. Distances between facing membranes were measured every 20 nm along the contact zone. A total of 90, 122 and 208 distances of contact made between tubes and spherical, hemi-spherical and angular-shaped vesicles were analyzed and depicted in the plot Figure 2C. In the plot shown in Figure 3C, a number of 52, 240, 491, 152 small vesicles have been analyzed for LPR 70, 175, 350 and 1400 lipid/VAP-A mol/mol, respectively.

#### Subtomogram averaging

Tubes containing PI(4) N-PH-FFAT and vesicles reconstituted with VAP-A were identified and segmented in Dynano (Castaño-díez et al., 2012; Castaño-Díez et al., 2017). The distance between tubes and vesicles was measured and particles were classified in classes corresponding to distance ranges of 0-5 nm, 5-10 nm, 10-15nm, 15-20 nm, 20-25 nm and 25-30 nm. Further analysis was performed with the most homogeneous class 20-25 nm. The full analysis is described in supplementary information (Figure S4). Briefly, subvolumes extracted from the center of the contact zone were aligned using a cylindrical mask covering both membranes and the contact region. A new round of alignment, in which the initial reference was low-pass-filtered to 48 Å, was run using a mask that covered the contact zone, where density corresponding to protein complexes was already observed. Features of the complex VAP-A with N-PH-FFAT were resolved but attempts to align them to higher resolution were unsuccessful, likely due to flexibility of the complexes. Thus, individual components N-PH-FFAT and VAP-A of the complex were analyzed separately. Angular search and shifts were gradually decreased while resolution was gradually increased by moving from bin4 to bin2 subvolumes and by moving the low-pass filter towards higher resolution. The resolution for VAP-A and of NPH-FFAT as determined by the FSC0.134 was 19.6 Å and 9.8 Å, respectively (Figure S4). Data processing are presented Table S1.

#### Structure prediction and modeling of proteins

Secondary structure predictions of human VAP-A (Q9P0L0) and N-PH-FFAT i.e. 1-408-OSBP have been performed using Phyres2, PSIPRED 4.0 (Buchan and Jones, 2019; Kelley et al., 2016).

3D models of 195-335 OSBP were generated using the homology method provided by Robetta, the protein structure prediction server (Yang et al., 2020) (Figure S5B).

#### Molecular Dynamics (MD) simulation of dimer model

The 3D model of 195-335 OSBP (Model 1, Figure S5B) generated from Robetta was duplicated to create two chains and positioned with on average 9Å distance between them with Pymol software (Schrödinger, 2010). MD simulations were performed with GROMACS 2019.4 (Abraham, M.J. et al., 2015) with the CHARMM36 force field (Huang, J., and MacKerell, 2013). The dimer was centered in the cubic box, at a minimum distance of 2.5 nm from box edges. Solvent molecules were added in the coordinate file. The TIP3P water model configuration was used (Jorgensen et al., 1983). Na and Cl ions were added to neutralize the simulation box and at a minimum concentration of 120 mM. The total number of atoms (protein + solvent + counterions) was 958,802.

Energy minimization was performed using the steepest descent minimization algorithm for the subsequent 50,000 steps. A step size of 0.01 nm was used during energy minimization. A cut-off distance of 1 nm was used for generating the neighbor list and this list was updated at every step. Long-range electrostatic interactions were calculated using the particle mesh Ewald summation methods (Darden, T., York, D., and Pedersen, 1983). Periodic boundary conditions were used. A short 100 ps NVT equilibration was performed. During equilibration, the protein molecule was restrained. All bonds were constrained by the LINear Constraints Solver (LINCS) constraint-algorithm (Hess, 2008). A short 100 ps NPT equilibration was performed similar to the NVT equilibration. During the production run, the V-rescale thermostat and Parrinello-Rahman barostat (Parrinello and Rahman, 1981) stabilized the temperature at 300 K and pressure at 1 bar, respectively. The simulations were performed for 300 ns and coordinates were saved every 100.0 ps. The MD analysis (distance between Nter/Cter) were performed using Gromacs utilities. Similar procedure was performed with the 3D model of 195-335 OSBP (Model 2, Figure S5B) without leading to a stable dimer.

#### Visualization

Rendering of 3D density maps is performed with UCSF Chimera (Goddard et al., 2007). Rigid-body docking of the crystallographic structure of the dimer of VAP-A MSP (PDB ID code 1Z90) and of the computed (195-335)-OSBP dimer into the cryo-EM density maps was done with Chimera.

## SUPPLEMENTARY INFORMATION

**Figure S1.**
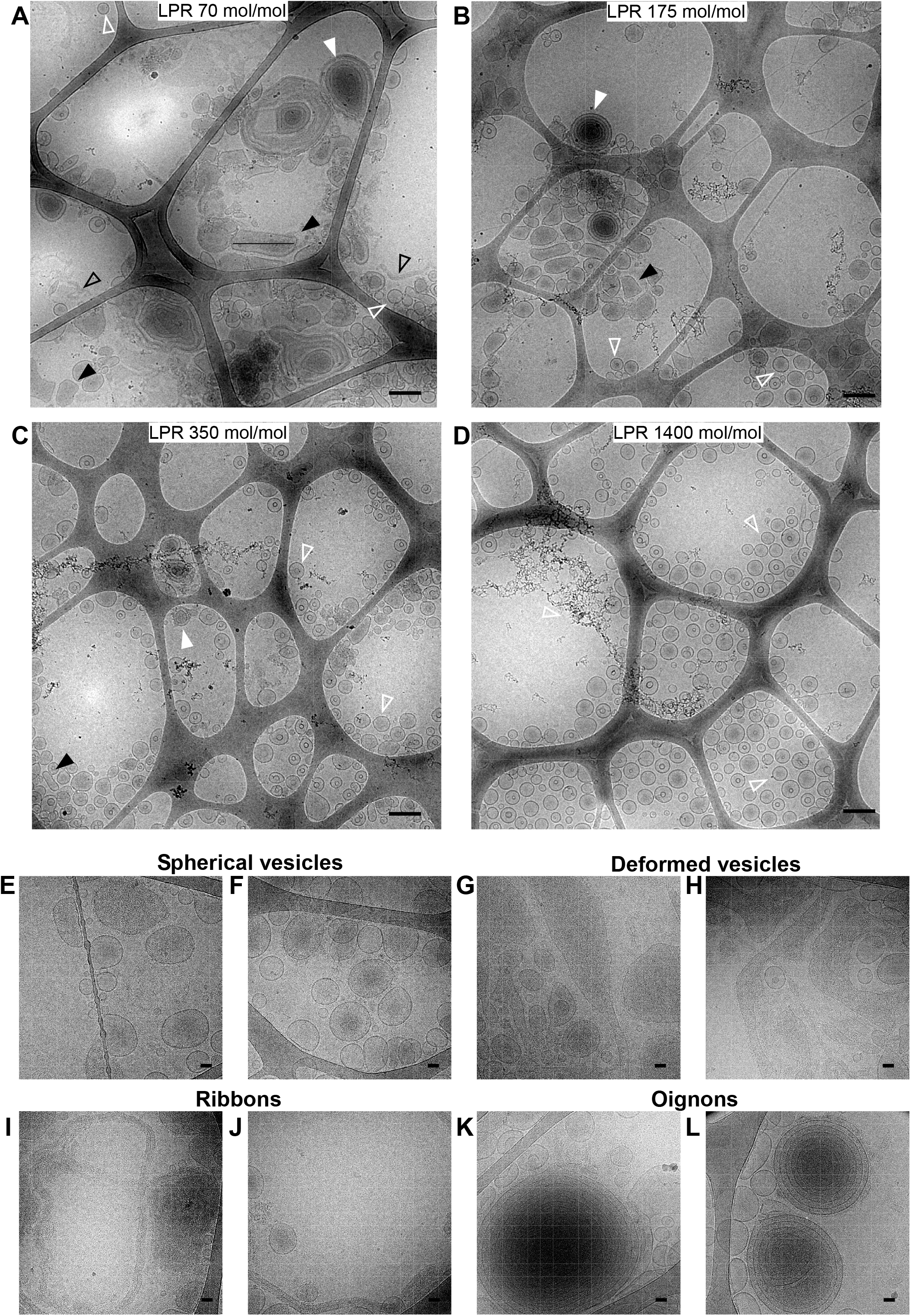
Cryo-EM images of reconstitution of VAP-A at different lipid/protein molar ratio. (A) Reconstitution of VAP-A at LPR 70, (B) 175, (C) 350, (D) 1400 mol/mol. Onions (full white arrow), deformed vesicles (full black arrow), ribbons (open black arrow), spherical liposomes (empty white arrow). Bars = 250 nm. (E-L). Different types of reconstituted membranes containing of VAP-A at LPR 70 and 175 mol/mol. (E, F) spherical and slightly deformed vesicles, (G, H) deformed vesicles, (I, J) ribbons, (K, L) onions. Bars: 50 nm.

**Figure S2.**
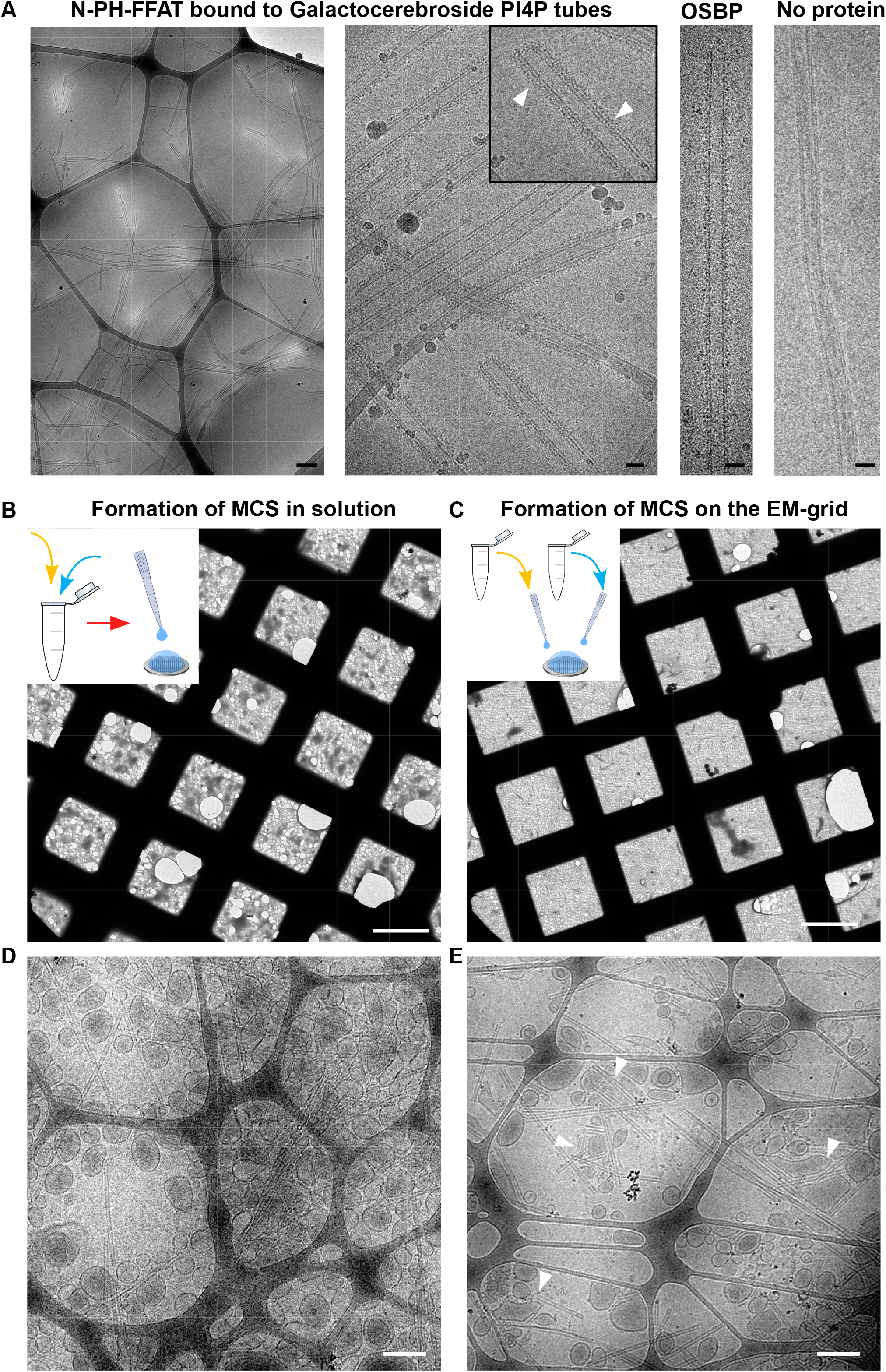
Formation of in vitro membrane contact site. (A) N-PH-FFAT or OSBP bound to galactocerebroside tubes doped with PI4P. Galactocerebroside tubes Galcer/EPC/PS/Pi4P (80/10/5/5) were mixed with N-PH-FFAT at 80 lipid/protein mol/mol. Tubes are covered with proteins that point to 6 nm from the membrane (white arrows Inset). Bars: 250 nm (left figure), 25 nm. (B-E). Formation of membrane contact site for cryo-EM on cryo-EM grids. (B, D) The mixture of VAP-A proteoliposomes and N-PH-FFAT-tubes in an eppendorf before freezing lead to a massive aggregation of material and thick ice. (C, E) Sequential addition of VAP-A proteoliposomes and N-PH-FFAT tubes on cryo-EM grids lead to the formation of membrane contact site (white arrow heads) suitable for cryoEM and cryo-ET. Bars: 50 μm in (B, C) and 250 nm in (D, E).

**Figure S3.**
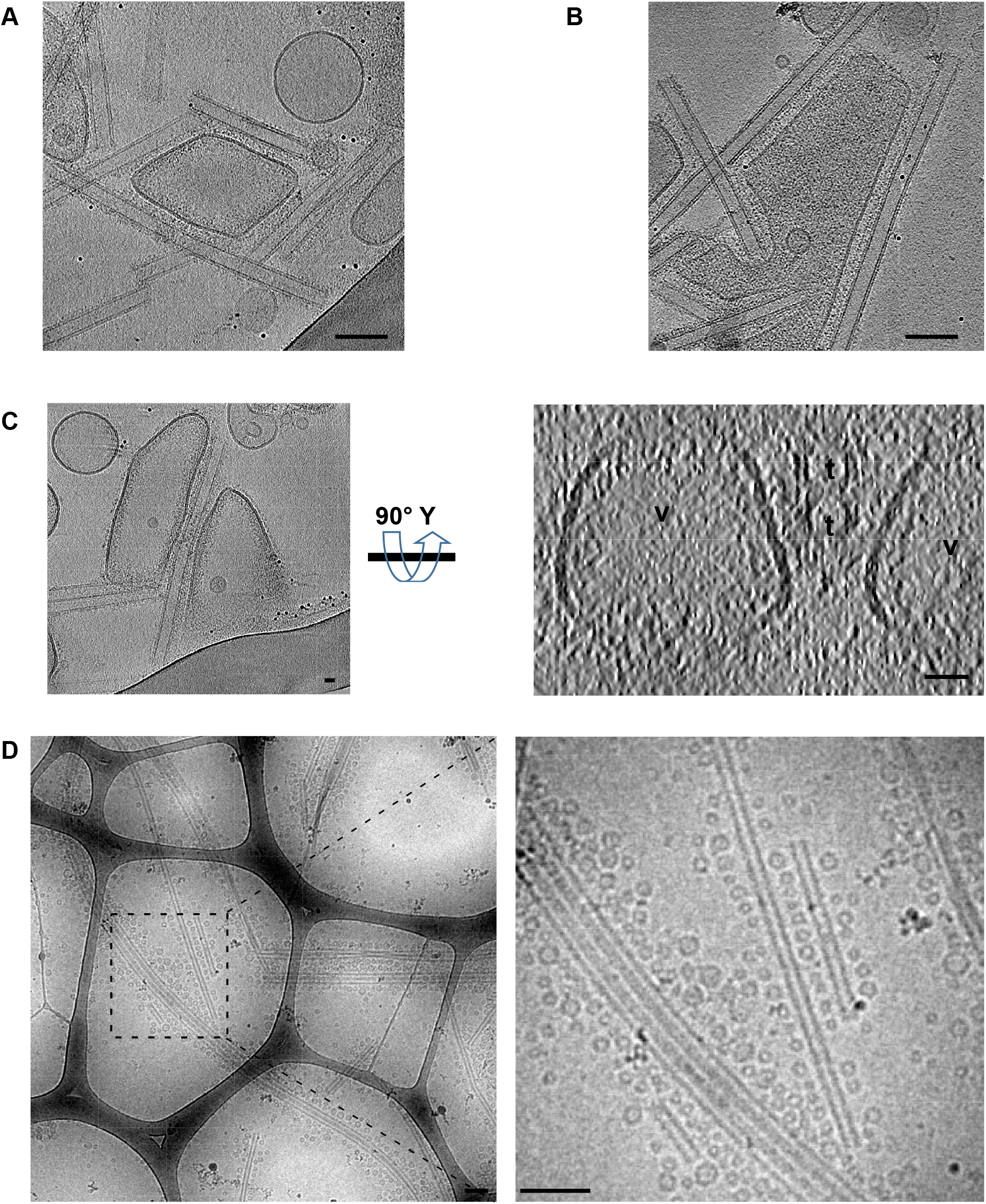
Membrane remodeling during formation of membrane contact sites. (A, B) Tomographic slices of two tilt series of VAP-A vesicles remodeled in contact with 4 tubes. In B, the large vesicle went around the tube but did not wrapped around. Bars: 100 nm (C) Tomographic slice of a tilt series in XY and YZ plans. In YZ plan, VAP-A vesicles (v) are flattened in contact to two close tubes (t). Bars: 25 nm (D) Low magnification and inset of MCS made with small vesicles of VAP-A reconstituted at LPR 175 mol/mol and N-PH-FFAT tubes. Bars: 100 nm.

**Figure S4.**
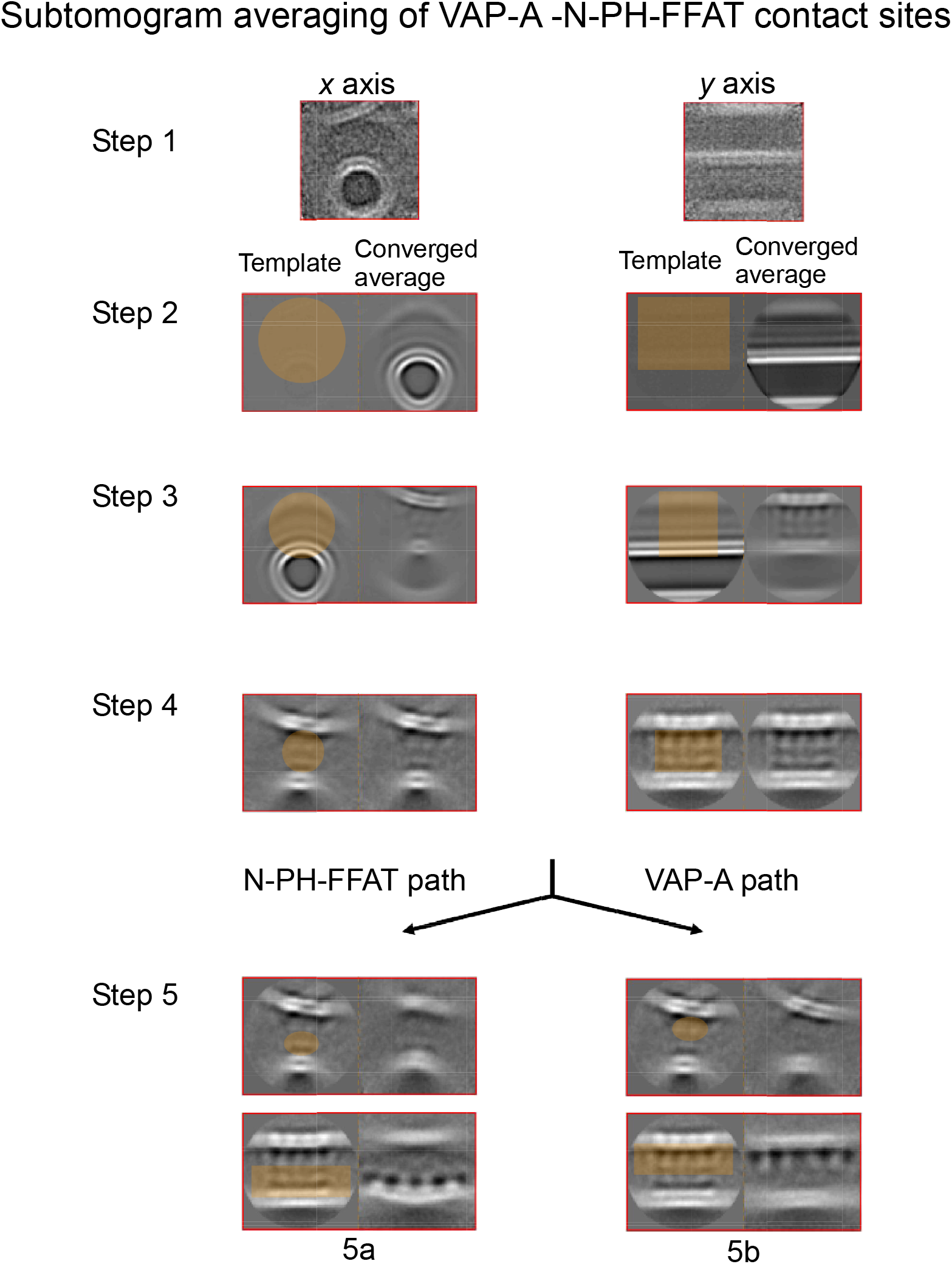
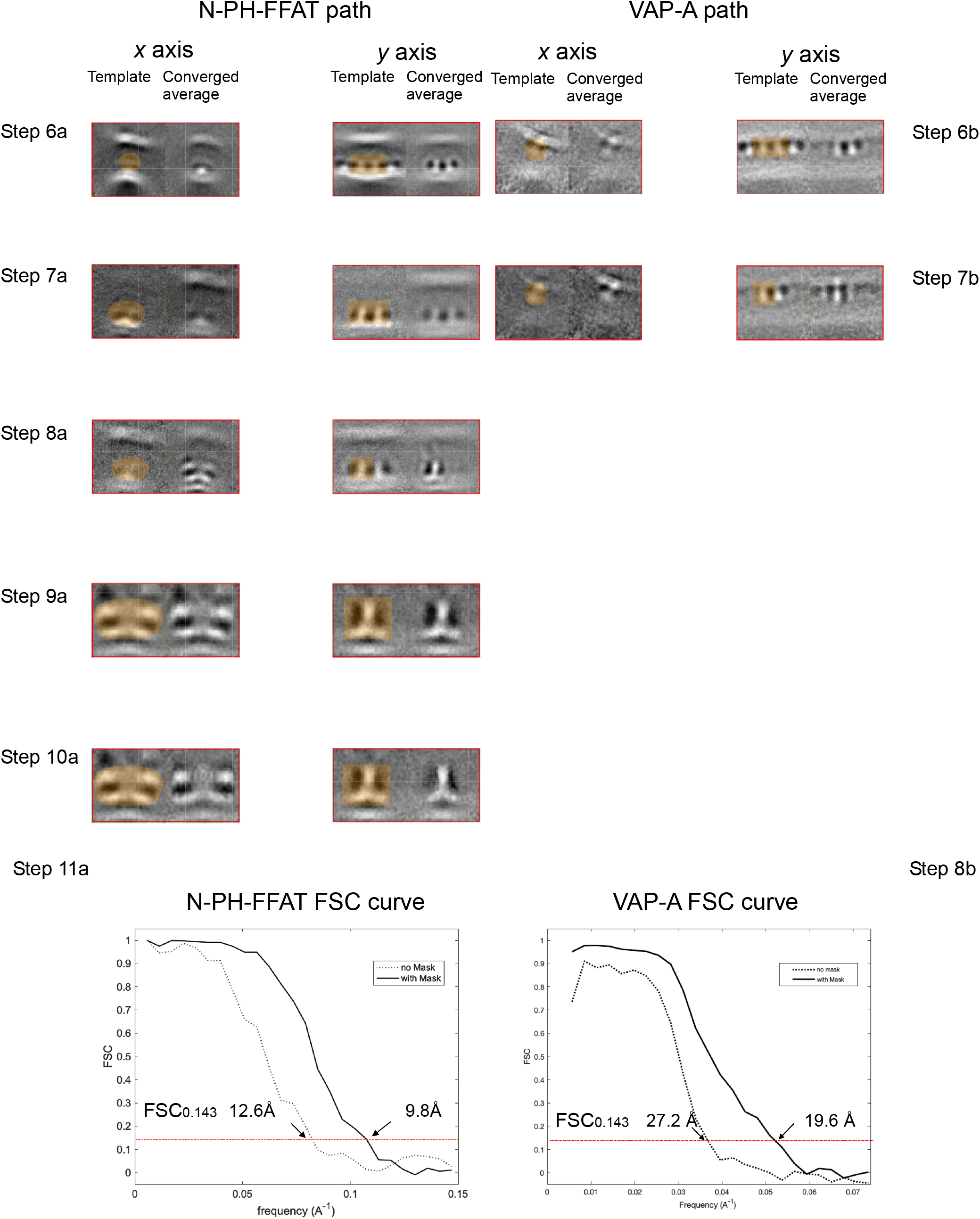
Detailed pipeline of subtomogram averaging. All steps of the following pipeline were performed under Dynamo (Castaño-díez et al., 2012; Castaño-Díez et al., 2017). **Step 1** During the first step the whole dataset composed of 63,348 oversampled subvolumes of 88^3^ voxels (59.8^3^ Å) belonging to class III (Figure 4B) were extracted from binned 4 (voxel size = 6.8 Å) tomograms, from which an initial model-free template was obtained by averaging a set of 2,000 randomly chosen subvolumes. **Step 2** Membranes from tubes and vesicles were used during the first rounds as references to align apposed membranes. A cylindrical mask covering half of the tube and the upper region corresponding to vesicles membrane was selected. Particles were binned 2X and the angular search was restricted to a cone range of 24° (steps of 6°) and to an in-plane range of 32° (steps of 8°). Particles were allowed to shift 2 voxels along the x, y and z axes. The low-pass filter was set to 40 Å. **Step 3** Once the membranes were aligned, two lines of density corresponding to the proteins involved in the contact were observed between the apposed membranes. During step 3 we reduced the size of the cylindric mask to align the region occupied by proteins. The angular search for binned 2 particles was set to a cone range of 16° (steps of 4°) and an in-plane range of 24° (steps of 6°). **Step 4** When proteins from both sides of the membrane were observed, the size of the aligned subvolumes was reduced to 72^3^ voxels (49^3^ Å), being the center of box that of the contact zone, i. e. the midpoint between the external layer of tube and vesicle membranes. An alignment with restricted parameters, no angular search (zero degrees) and shifts of 1 voxel along the axis *x, y* and *z*, was performed as a control. Particle size was set to 36 (bin 2) and low-band pass to 40 Å. **Step 5** N-PH-FFAT bound to PI4P contained in tube membranes and VAP-A reconstituted in vesicles were aligned following independent pipelines. The goal of this step was to achieve a better alignment of the individual components of the contact zone. A mask covering either N-PH-FFAT (step 5a) or VAP-A (step 5b) was chosen. Angular search was restricted to a cone range of 4° (increments of 1°) and an in-plane range of 4° (increments of 1°). Shifts were set to 8, 2 and 2 along *x, y* and *z*, where *x* corresponds to the tube axis. Particle size and low-band pass were kept as in last step. **Step 6 a,b** As a result of the alignment described in step 5, four subunits of both N-PH-FFAT and VAP-A were clearly distinguished in the contact region of the aligned subvolumes. The goal of this step was to align the two subunits located in the central region of the box. Angular search parameters were kept to 4 ° (steps of 1°) for both cone and in-plane ranges. Shifts were also kept to 8, 2, 2, and particle separation was set to 8 voxels (5.4 Å) to decrease the number of subvolumes and select those with the best correlation coefficient (CC). Particle size and low-band pass were kept as before, i. e. 36 (bin 2) and 40 Å, respectively. **Step 7 a,b** Aligned subvolumes with dimensions of 104^3^ voxels (35.4^3^ Å) were extracted from binned 2 (voxel size = 3.4 Å) tomograms. An alignment with restricted parameters, zero degrees for angular search and shifts of 1, 1, 1 voxel, was performed as a control to confirm that the new extraction didn’t modified the result observed in step 6b. Particle size was set to 52 (bin 2) and low-band pass to 25.2 Å. The alignment mask covered the lower central region of the box. The following steps were only performed over the N-PH-FFAT path. **Step 8a** The alignment mask was restricted to one of the two subunits of N-PH-FFAT observed in the template. Angular search was set to 4° (increments of 1°) for both, cone and in-plane range. Shifts were set to 16, 2 and 2 voxels. Both membranes, that from PI4P tubes and that from VAP-A containing vesicles are still observed in the resulting average. Particle size and low-band pass were set to 36 voxels (bin 2) and 25.2 Å, respectively. **Step 9a** Aligned subvolumes were *subboxed* to reduce the box size to 52^3^ voxels (17.7^3^ Å) and to shift the center of the box towards the aligned subunit of N-PH-FFAT. A restricted alignment was run to confirm that subvolumes were properly *subboxed*. Symmetry was set to c2 based on the evident two-fold symmetry observed, the symmetry test performed under Dynamo and on biochemical data that have shown that N-PH-FFAT is a dimer (Jamecna *et al*. 2019). **Step 10a** The last step consisted of an adaptive band-pass filter alignment (FSC set to 0.143). Initial parameters of the angular search were set to a cone range of 16° (increments of 4°) and in-plane range of 180° (increments of 10°) while shifts were set 8, 6, 6 voxels along the *x*, *y* and *z* axes. Symmetry was kept as c2. Both, the angular search and shifts were gradually decreased, and the band-pass filter moved towards higher resolution. **Step 11a, 8b** The final converged average consisted on 15,597 and 6,948 subvolumes for N-PH-FFAT and VAP-A respectively, and their respective FSC were calculated over the two half-averages obtained upon randomly splitting the subvolumes in two datasets.

**Figure S5.**
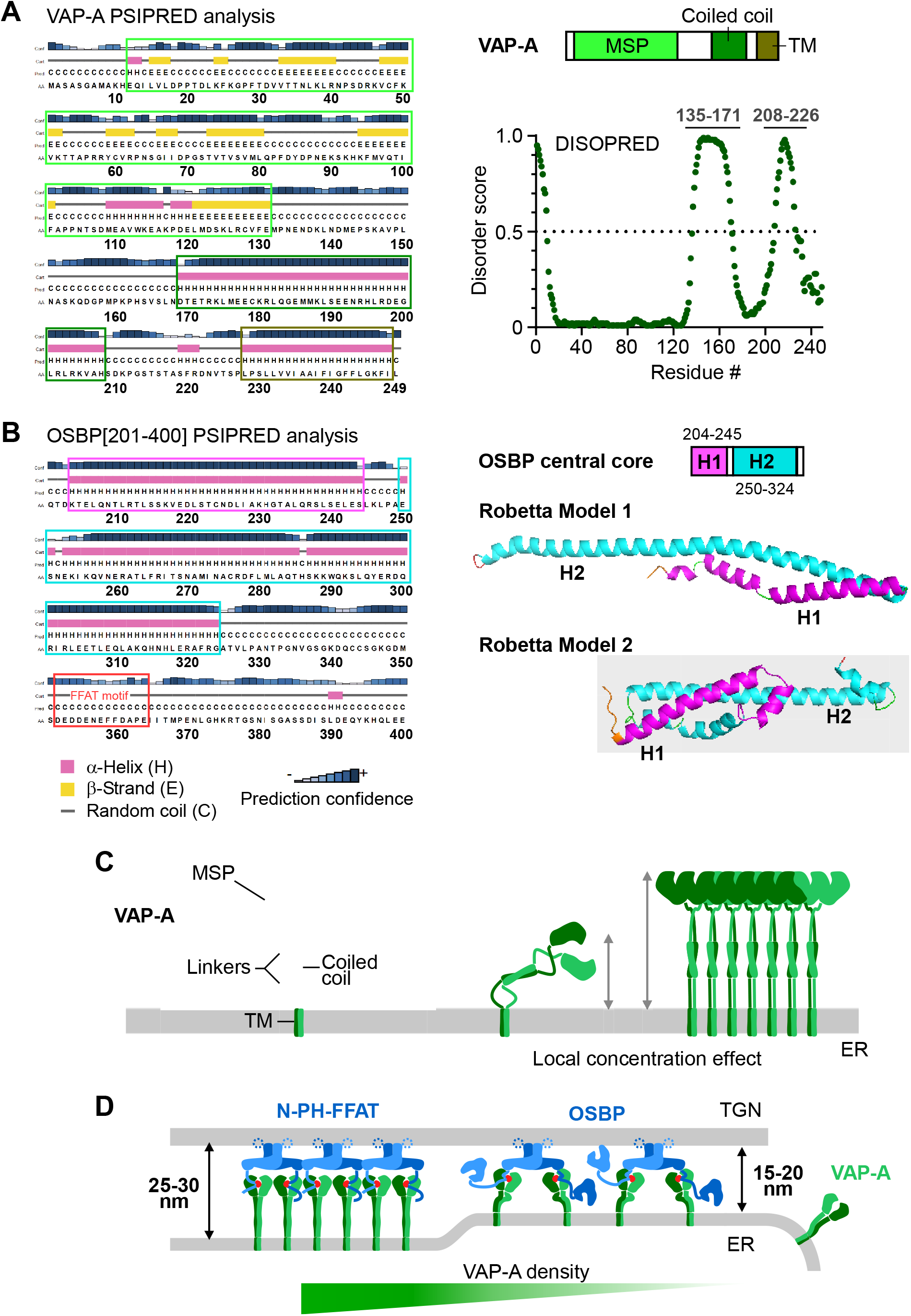
Prediction and model of VAP-A and OSBP. (A) Secondary structure prediction of human VAP-A by PSIPred. VAP-A is made of the MSP domain (light-green square), a putative CC region (green square), a transmembrane helix (brown square) and two unstructured linkers SL1 (aa135-171) and SL2 (aa208-226) that flank the CC region. Disordered regions predicted by Disopred program (B) Secondary structure prediction of human central core OSBP (aa201-400) by PSIPred program. OSBP (aa201-400) is helix H1 (aa204-245) and helix H2 (aa249-325) and a disordered region 326-408 that contains the FFAT motif. Models of central core OSBP computed by Robetta (C) Schematic representation of VAP-A organization at the ER membrane. The disordered linkers of VAP-A allow VAP-A to explore a large space to bind its interactors. When concentrated in a cluster VAP-A can extend to 17 nm from ER membrane. (D) Organization of VAP-A in ER-Golgi MCS depending on its local concentration. VAP-A concentrate in flat membrane region of MCS and adapt its concentration to its interactors. Fewer molecules of VAP-A are engaged in MCS containing OSBP with bulky ORD domains than with N-PH-FFAT. This results in shorter intermembrane distances in MCSs containing OSBP than N-PH-FFAT.

Movie S1: Tomographic volume of VAP-A-N-PH-FFAT membrane contact sites and annotated segmentation of the tomographic data. Related to Figure 2D. White: large and flat VAP-A vesicle, small spherical VAP-A vesicle. Blue/black two lipidic tubes with bound N-PH-FFAT.

Movie S2: Tomographic volume of VAP-A-N-PH-FFAT membrane contact sites and annotated segmentation of the tomographic data. Related to Figure 2D. White: ribbons like VAP-A membrane. Black lipidic tubes with bound N-PH-FFAT.

Movie S3: Tomographic volume of VAP-A-OSBP membrane contact sites and annotated segmentation of the tomographic data. Related to Figure 2D. White: large and flat VAP-A vesicle, small spherical VAP-A vesicle. Blue/black two lipidic tubes with bound OSBP.

Movie S4: Tomographic volume of VAP-A-N-PH-FFAT membrane contact sites. Two flat vesicles spread along a tube and are connected by a short tongue. Related to Figure 3A.

